# A Library of Phosphoproteomic and Chromatin Signatures for Characterizing Cellular Responses to Drug Perturbations

**DOI:** 10.1101/185918

**Authors:** Lev Litichevskiy, Ryan Peckner, Jennifer G. Abelin, Jacob K. Asiedu, Amanda L. Creech, John F. Davis, Desiree Davison, Caitlin M. Dunning, Jarrett D. Egertson, Shawn Egri, Joshua Gould, Tak Ko, Sarah A. Johnson, David L. Lahr, Daniel Lam, Zihan Liu, Nicholas J. Lyons, Xiaodong Lu, Brendan X. MacLean, Alison E. Mungenast, Adam Officer, Ted E. Natoli, Malvina Papanastasiou, Jinal Patel, Vagisha Sharma, Courtney Toder, Andrew A. Tubelli, Jennie Z. Young, Steven A. Carr, Todd R. Golub, Aravind Subramanian, Michael J. MacCoss, Li-Huei Tsai, Jacob D. Jaffe

**Affiliations:** The Broad Institute, 415 Main St., Cambridge, MA 02142 USA; Department of Brain and Cognitive Sciences, Picower Institute for Learning and Memory, Massachusetts Institute of Technology, 77 Massachusetts Avenue, Cambridge, MA 02139, USA.; University of Washington, Department of Genome Sciences, 3720 15th Ave NE, Seattle WA 98195, USA

**Keywords:** Mass spectrometry, proteomics, proteomic profiling, drug discovery, signaling, chromatin, epigenetics, mechanism of action

## Abstract

Though the added value of proteomic measurements to gene expression profiling has been demonstrated, profiling of gene expression on its own remains the dominant means of understanding cellular responses to perturbation. Direct protein measurements are typically limited due to issues of cost and scale; however, the recent development of high-throughput, targeted sentinel mass spectrometry assays provides an opportunity for proteomics to contribute at a meaningful scale in high-value areas for drug development. To demonstrate the feasibility of a systematic and comprehensive library of perturbational proteomic signatures, we profiled 90 drugs (in triplicate) in six cell lines using two different proteomic assays — one measuring global changes of epigenetic marks on histone proteins and another measuring a set of peptides reporting on the phosphoproteome — for a total of more than 3,400 samples. This effort represents a first-of-its-kind resource for proteomics. The majority of tested drugs generated reproducible responses in both phosphosignaling and chromatin states, but we observed differences in the responses that were cell line-and assay-specific. We formalized the process of comparing response signatures within the data using a concept called connectivity, which enabled us to integrate data across cell types and assays. Furthermore, it facilitated incorporation of transcriptional signatures. Consistent connectivity among cell types revealed cellular responses that transcended cell-specific effects, while consistent connectivity among assays revealed unexpected associations between drugs that were confirmed by experimental follow-up. We further demonstrated how the resource could be leveraged against public domain external datasets to recognize therapeutic hypotheses that are consistent with ongoing clinical trials for the treatment of multiple myeloma and acute lymphocytic leukemia (ALL). These data are available for download via the Gene Expression Omnibus (accession GSE101406), and web apps for interacting with this resource are available at https://clue.io/proteomics.

**Highlights:**

- First-of-its-kind public resource of proteomic responses to systematically administered perturbagens
- Direct proteomic profiling of phosphosignaling and chromatin states in cells for 90 drugs in six different cell lines
- Extends Connectivity Map concept to proteomic data for integration with transcriptional data
- Enables recognition of unexpected, cell type-specific activities and potential translational therapeutic opportunities

## Introduction

Molecular profiling technologies have enabled tremendous advances in biomedicine ranging from basic mechanistic insights to active guidance of therapeutic choices in precision medicine. For example, a gene expression signature (PAM50) now constitutes one of the major diagnostic classifiers for breast cancer (Parker et al., 2009). Gene expression profiling has long held sway as the technology of choice for generating systematic, comprehensive data sets of sufficient size to power statistical analyses. Early landmark studies (Alizadeh et al., 2000; Bittner et al., 2000; Clark et al., 2000; Golub et al., 1999; Perou et al., 2000) demonstrated the power of molecular profiling, paved the way for its acceptance into the mainstream, and ultimately drove costs down and technology forward. It has even been proposed that gene expression profiling could serve as the “‘universal language’ with which to describe cellular responses,” from which the concept of the Connectivity Map — linking drugs, genes, and phenotypes through expression profiles — has emerged (Lamb et al., 2006; Subramanian et al., 2017).

Yet it is known that mRNA levels alone do not fully capture cell state, and the vocabulary of this universal language may need to be extended to describe some of the critical functions of cellular responses. Early studies recognized apparent discordance between mRNA and protein levels on a large scale (Greenbaum et al., 2003), and current studies suggest there is correlation coefficient of about ~0.5 between mRNA and protein levels (Mertins et al., 2016). More recently, Li and colleagues have shown that phosphorylation events show low correlation with mRNA levels from their corresponding genes (Li et al., 2017), and therefore phosphoproteomic data are likely to add value to gene expression measurements. These observations underscore the importance of exploring complementary readouts to gene expression profiling.

Alternative profiling methodologies could potentially fill the gaps in gene expression profiling by measuring analytes that cannot be detected via nucleic acid, reporting on biological processes with time-scales distinct from changes in gene expression, being scalable to prosecute systematic studies of sufficient size to power discovery, and yet remaining cost-effective enough to deploy on a routine basis. Further, it would be highly desirable for molecular profiling assays to directly report on cellular processes affected by novel drug candidates’ primary modes of action, which usually involve inhibition of protein activity rather than the achievement of a particular transcriptional state. Two emerging classes might particularly benefit from such directed assays: 1) targeted kinase inhibitors, a class of drugs that is rapidly expanding (Wu et al., 2015), and 2) inhibitors of chromatin-modifying enzymes and sensors of chromatin state, which have emerged as exciting new therapeutic modalities (Dawson et al., 2012; Kelly et al., 2010). While these new therapies are extremely promising, deeper insight is required to fully understand their cellular effects in their intended biological contexts, as well as their possible off-target effects in a system-wide manner. At the same time, many established protein-targeting drugs lack clear mechanistic insight and may harbor unexpected phosphosignaling and epigenetic activities that would be useful in repurposing efforts (Gupta et al., 2013; Singhal et al., 1999).

To fill these needs, we set out to develop a reference resource of proteomic signatures elicited in response to drugs, specifically monitoring phosphosignaling and chromatin state, two key areas for therapeutic development. Unlike recent proteomic resources characterizing the proteomic *states* associated with genetic variation in tumors and cell lines (Li et al., 2017; Mertins et al., 2016), this resource characterizes the proteomic *responses* to systematic application of drug perturbagens. Here, we describe the creation and validation of a pilot library containing signatures from our reduced-representation phosphoproteomic assay (Abelin et al., 2016) and our global chromatin profiling assay (Creech et al., 2015). These proteomic assays are highly standardized and rigorously quantitative, yet automated and compact enough to achieve a relatively high throughput and reasonable scale. We have systematically profiled a wide variety of approved drugs and tool compounds across a range of biological models, including cancer and neurodevelopment. We show that proteomic profiling yields reproducible signatures that enable recognition of mechanisms of action and cell type-specific effects. The relatively large scale of the data (compared to other proteomic perturbation studies) allows for a principled query of proteomic signatures that, for the first time, allow us to extend the Connectivity Map concept (Lamb et al., 2006; Subramanian et al., 2017) to proteomics data and easily integrate proteomic with transcriptomic data. We show that the resource itself contains a wealth of information about drug activity in cells, and also that the resource can be leveraged for analysis of external data to recognize potential therapeutic opportunities. This data resource contributes to the NIH Library of Integrated Network-based Cellular Signatures (LINCS) program, and represents a first-in-class queryable proteomics resource of over 3,400 profiles. Importantly, the resource continues to grow beyond its current size with longitudinal support from LINCS and opens the door to both utilization by and contributions from the community at large.

## Results

### Structure and scope of the proteomic signature resource

We created a library of proteomic perturbational signatures containing more than 3,400 samples (Figure 1A). In this initial dataset, we profiled 90 small molecules spanning a variety of mechanisms of action (MoAs) with focused subsets of kinase inhibitors and epigenetically active compounds (Table S1). We utilized five widely studied cancer cell models (representing breast, lung, pancreatic, prostate, and skin lineages) and one neurodevelopmental cell model — neural progenitor cells (NPCs) — in order to test non-cancer models. Samples were profiled with a reduced-representation phosphoproteomic assay (P100) and a global chromatin profiling (GCP) assay, both of which are liquid chromatography-mass spectrometry (LC-MS)-based and previously described (Abelin et al., 2016; Creech et al., 2015). The analytes measured by P100 are ~100 phosphorylated peptides from cellular proteins, and the analytes measured by GCP are ~60 post-translationally modified peptides from histones (e.g. methylated, acetylated, and combinations thereof). The analytes measured by P100 serve as a reduced representation of the phosphoproteome and act as early sentinels of bioactivity of a diverse set of signaling pathways and drug mechanisms. The analytes measured by GCP include nearly every well-studied post-translational modification on the core nucleosomal histone proteins. These modifications convey epigenetic information in cells and their dysregulation is associated with a wide range of diseases (Araf et al., 2016; Aumann and Abdel-Wahab, 2014; Gräff and Mansuy, 2009; Jaffe et al., 2013; Ntziachristos et al., 2016; Peña et al., 2014).

**Figure 1.**
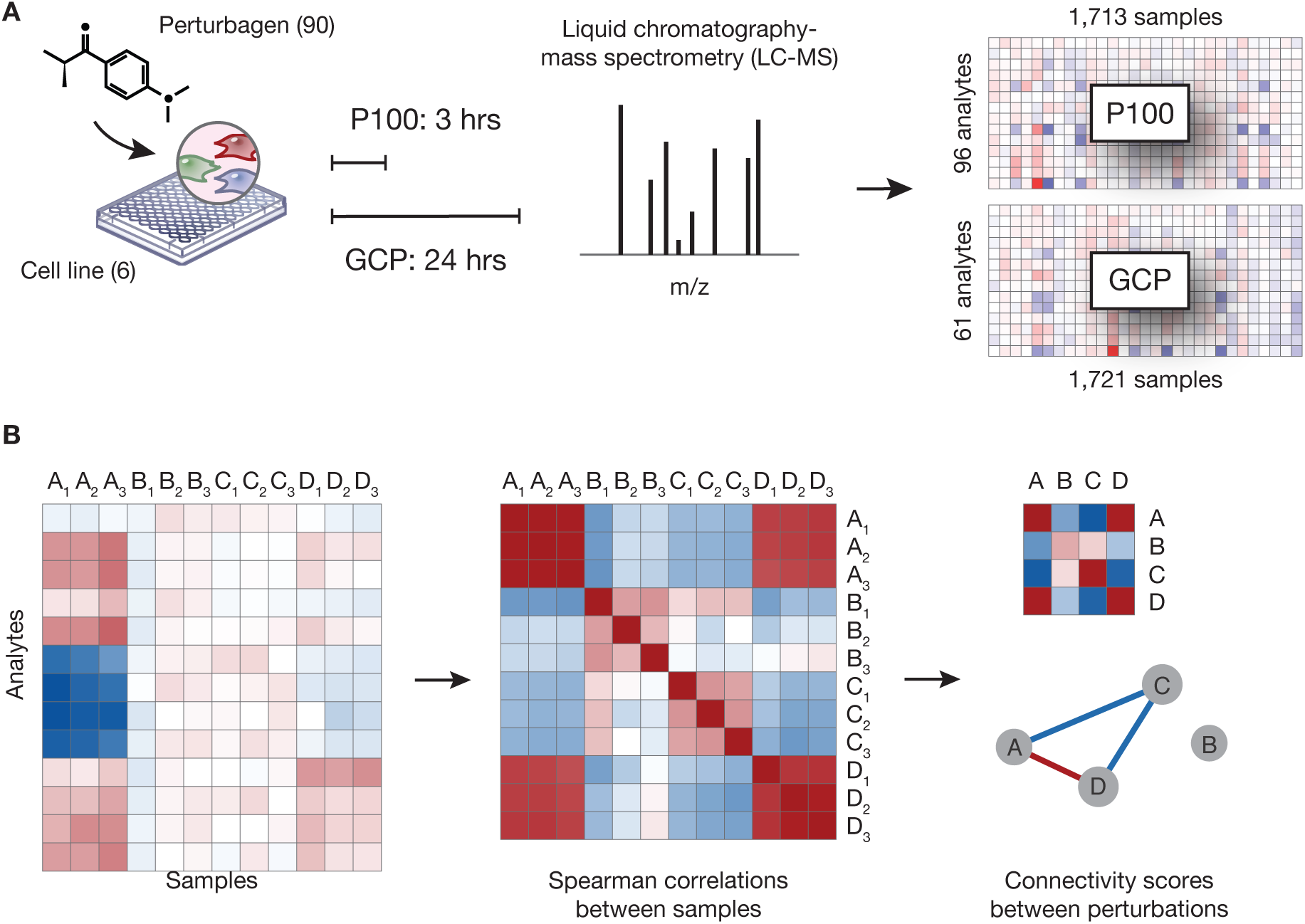
Overview of experiment and computational framework. (A) Experimental design. Cells were treated with one of 90 small-molecule perturbagens with a minimum of three biological replicates. After 3 (P100) or 24 (GCP) hours of exposure to treatment, cells were lysed and profiled in each of the two assays. After quality control filtering, over 3,400 individual profiles comprised the resource. (B) Computational framework. Each sample is represented as a profile of analyte measurements. Spearman correlations are computed between all profiles within a cell line. Finally, we compute connectivity scores by comparing the observed correlations to a background of correlations. Computing connectivity collapses replicates. Connectivity maps may be represented as matrices or networks. See also Figure S1.

We adopted guiding principles governing the generation of the resource (Supplemental Note 1). First, we chose time scales that were relevant for the biological processes covered by the assays: 3 hours after drug treatment for phosphoproteomic profiling and 24 hours for chromatin profiling. Second, we chose treatment concentrations consistent with known bioavailability levels in humans where possible. Sample preparation for each assay was highly automated, which allowed for preparation and analysis of 96 samples per batch. The final output of profiling was matrix data — analytes in the rows and samples in the columns (Figure 1A, far right) — that can be readily analyzed by modern computational techniques. All of our data are publically available (Table 1). Extensive metadata concerning the treatment parameters and analyte identities were also embedded in the matrices, according to the standards set forth by the Library of Integrated Network-based Cellular Signatures (LINCS) consortium (http://www.lincsproject.org). Beyond the analysis presented here, we are committed to expansion of the resource and have already released primary signature data for genetic knockdowns by CRISPR, time course of drug response in MCF10A cells, and drug responses in cardiovascular cell models via our Panorama repository.

**Table 1.** Web-accessible persistent data locations.

| Type of data | Location | URL |
| --- | --- | --- |
| Extracted ion chromatogram MS data | Panorama Web | https://panoramaweb.org/labkey/LINCS.url |
| Signatures | Gene Expression Omnibus (GEO), accession GSE101406 | https://www.ncbi.nlm.nih.gov/geo/query/acc.cgi?acc=GSE101406 |
| Connectivities | CLUE web apps | https://clue.io/proteomics |

### Universal connectivity framework for data processing and integration

Mass spectrometry data are summarized by Skyline software (MacLean et al., 2010) to yield a quantification for each analyte of the amount of peptide that was detected relative to a spiked-in synthetic control. Further data processing is performed by the Proteomics Signature Pipeline (https://github.com/cmap/psp). Briefly, the data are log_2_-transformed, analytes and samples with an excess of missing data are filtered out, a constant offset is applied to each sample to bring its range of values to the same scale as the other samples on a plate, and analytes are median normalized (see STAR Methods for a detailed description of the processing pipeline). All data presented in this study, at multiple levels of processing (Figure S1A), have been consolidated under a stable Gene Expression Omnibus record (GEO, accession GSE101406).

We adopted the concept of connectivity as a consistent, assay-independent means of identifying two or more drugs that elicit similar responses in cells (Lamb et al., 2006). It enables us to ask the question “which compounds look alike in our dataset?” without focusing on the biological meanings of the individual analytes measured. For example, by recognizing a connection between a drug of known mechanism and one of unknown mechanism, we can formulate a hypothesis about the unknown drug.

An important consequence of connectivity is the ability to integrate data *across* assays. Because our assays measure different analytes, we are unable to directly compare perturbations’ signatures. Instead, we compare their *connectivities*. In other words, we ask “does drug X have the same connections to other drugs in GCP as it does in P100?” This framework of connectivity allows us to quantitatively compare perturbations both within an assay and across assays, and connectivity scores have the same range regardless of assay. By taking a perturbation-centric approach rather than an analyte-or gene-centric approach, we can easily combine data from different assays using simple matrix operations (see results pertaining to assay comparison and data integration, below).

We compute connectivity in two steps (Figure 1B, see STAR Methods for a detailed description). First, we compare samples to each other using Spearman correlation, which considers entire signatures as the basis for comparison, rather than focusing on a limited number of up-or down-regulated analytes. Because correlation is sensitive to the number of analytes in a signature and our assays measure different numbers of analytes, we cannot directly compare correlations. Therefore, we convert correlations to *connectivity scores* by comparing observed correlations to a background distribution of correlations. In the process, we collapse replicates corresponding to the same perturbation. A connectivity score of 1 indicates that two perturbations are more similar to each other than all other pairs of perturbations, 0 indicates that their similarity is unexceptional, and −1 indicates that they are less similar to each other than all other pairs of perturbations (Figure S1B-D). Drugs with positive connectivity are highly likely to elicit the same underlying cellular *state* as reported by the original assay, with phosphopeptides as a proxy for signaling state in P100 and chromatin modifications as a proxy for epigenetic state in GCP). Drugs with negative connectivity likely elicit very different states with anti-correlated profiles.

Connectivity scores can be visualized using heatmaps or network views (Figure 1B, far right), and subjected to further quantitative summarization. Heatmaps can be manipulated as any other form of matrix data. For example, one can visualize the strongest connections to an *individual* compound by sorting a single column. One can visualize the strongest connections to a *set* of compounds (e.g. drugs corresponding to the same MoA) by summarizing their individual connectivities (e.g. by median) and sorting by this summary value. Complete matrices of proteomic connectivity data can be browsed and interactively manipulated at https://clue.io/proteomics. Network views are directed or undirected graphs representing perturbations as nodes and connectivity scores above a user-defined threshold as edges. Unlike heatmaps, network views make it easy to see different modules of biology at a glance. Automated methods for generating network views and input files suitable for Cytoscape visualization are provided as part of the Proteomics Signature Pipeline (see STAR methods) (Shannon et al., 2003).

### Proteomic assays produce reproducible signatures that capture diverse cellular responses

As a requisite first step, we assessed the technical quality of our datasets by quantifying replicate reproducibility. We compared the global distributions of replicate correlations to non-replicate (Spearman) correlations (Figure 2A). The distributions were significantly different according to a two-sample KS test (GCP: *D* = 0.63, *p* < 10^−15^; P100: *D* = 0.68, *p* < 10^−15^), indicating good separation of replicate correlations from non-replicate correlations. Compounds that lack bioactivity in a given system would not be expected to have high replicate reproducibility, and thus we expect some perturbations to behave poorly by this metric *a priori*.

**Figure 2.**
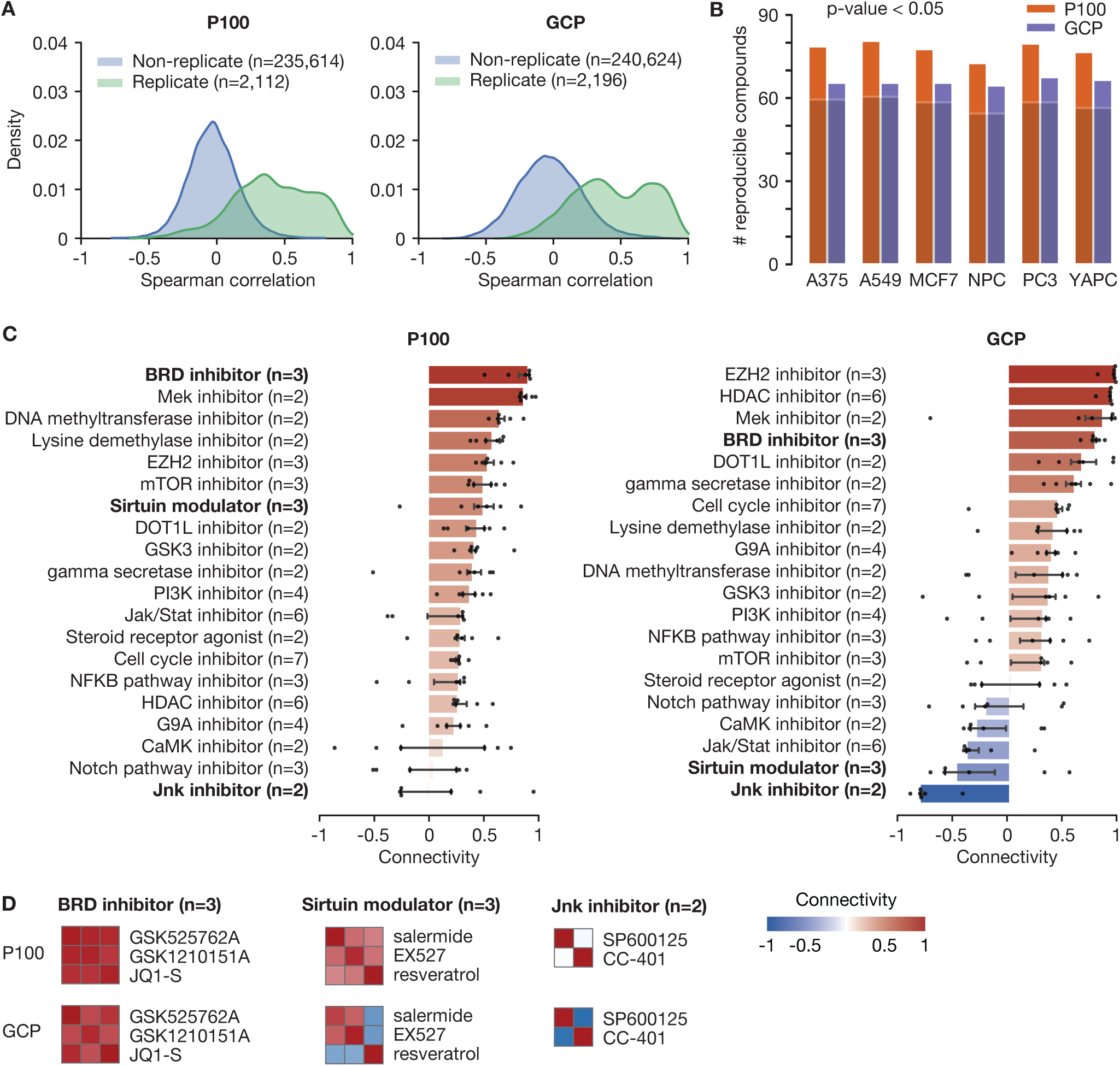
Majority of compounds are reproducible in both assays, but assays show different sensitivities to MoA classes. (A) Distributions of all Spearman correlations among replicates (green) and among non-replicates (blue). (B) Bar chart showing the number of compounds considered reproducible in each cell line-assay combination. A compound was considered reproducible if the correlations among its replicates were significantly higher (p-value < 0.05) than the correlations among randomly chosen samples. The shaded component indicates the overlap of reproducible compounds between GCP and P100. (C) Bar charts showing the median connectivity of compounds annotated with the same mechanism of action (MoA). The extent of each bar is the median of six cell-specific median connectivities; error bars represent the 25th and 75th percentiles. (D) Heatmaps of the connectivity among compounds belonging to the BRD inhibitor, sirtuin modulator, and JNK inhibitor MoA classes. The BRD inhibitor class shows high connectivity in both P100 and GCP; the sirtuin modulator class shows high connectivity in P100 but low connectivity in GCP; and the JNK inhibitor class shows low connectivity in both P100 and GCP. Each square is the median of six cell-specific connectivity scores. The labels of the matrices are symmetric; that is, the columns (left to right) have the same annotations as the rows (top to bottom). Color scale applies to both panels C and D. See also Figure S2.

Next, we asked how many individual perturbagens were reproducible in each cell line. A perturbagen was considered reproducible if the median of pairwise correlations between its replicates was significantly higher than the median of pairwise correlations between randomly chosen samples (see STAR Methods for a detailed description of the algorithm). At a *p*-value threshold of 0.05, we found that in each cell line in both assays, at least 64 out of the 90 compounds profiled (average = 71.2 compounds, or 79.1%) were reproducible (Figure 2B). Furthermore, in each cell line, at least 54 compounds (60%) were reproducible in both assays (shaded portion), indicating that a compound was likely to be reproducible in both assays if it was reproducible in one. These results are in line with the fraction of small-molecules determined to be bioactive using other profiling modalities: 68.3% for Cell Painting morphological profiling (Wawer et al., 2014) and 38% for L1000 gene expression profiling (Subramanian et al., 2017).

In each cell line, more compounds were reproducible in P100 than in GCP (mean difference = 11.6 compounds). Part, but not all, of this observation is explained by the larger feature space of P100 (Supplemental Note 2, Figure S2). In addition, compounds that were reproducible in P100 but not GCP in at least four cell lines (gossypetin, rolipram, olaparib, TBB, tacrolimus, and everolimus) tended to be annotated with kinase-directed activities, perhaps explaining the remainder of compounds that were only reproducible in the phosphoproteomic assay. Finally, we note that replicate reproducibility was similar between NPCs and the cancer cell models. This observation emphasizes that both assays work as well in unusual cellular contexts, such as a neurodevelopmental cell line, as in the more typical cancer cell models.

### Using proteomic connectivities to detect and refine mechanisms of action

Having validated the technical quality of our datasets, we sought to investigate how well various annotated MoAs were detected by each assay. For this analysis, we define intra-class connectivity as the median connectivity score among compounds belonging to the same MoA (Figure 2C; see Table S1 for compound annotations). In both assays, the majority of MoA classes had positive intra-class connectivities. Several MoA classes, such as bromodomain (BRD) inhibitors and MEK inhibitors, had high connectivity in both assays (Figure 2D, left).

Other MoA classes, such as JNK inhibitors, had low — in fact, negative — intra-class connectivity in both assays (Figure 2D, right). In the case of the two JNK inhibitors that we profiled, both compounds had strong self-connectivity (indicating high replicate reproducibility and therefore a definitive signal of each in both assays) but failed to connect to each other. We posit that this lack of connectivity reflects either mis-annotation of the compounds in literature, or, more likely, significant but distinct off-target effects in one or both at the doses tested. Indeed, a screen of 20 kinase inhibitors highlighted one of our two JNK inhibitors — SP600125 — as having very poor separation between its affinity for intended targets and off-targets, indicating that this drug has substantial off-target effects (Fabian et al., 2005). Our data show that the next-best connected compound in our dataset to SP600125 in P100 is the JAK2 inhibitor TG101348. This example illustrates the power of using unbiased assays and our connectivity framework to recognize annotation ambiguities and suggest alternative hypotheses about MoA.

Connectivity analysis can provide refinement to ambiguous MoA classes in an assay-specific manner. We included three compounds thought to modulate the activity of the sirtuin class of histone deacetylases: resveratrol, EX527, and salermide. Resveratrol is typically annotated in literature as a sirtuin activator (Wendling et al., 2013), but it has also been argued that resveratrol does not directly act on sirtuins at all (Beher et al., 2009; Pacholec et al., 2010). Because of this ambiguity, we grouped resveratrol and the two known sirtuin inhibitors into a single MoA class. In our analysis, this class had positive connectivity in P100 (0.49) but negative connectivity in GCP (-0.47). Upon investigation, we discovered that the negative connectivity in GCP is explained by resveratrol having strong negative connections to the other two compounds in the class, EX527 and salermide (Figure 2D, middle). The negative connectivity in GCP between resveratrol and the sirtuin inhibitors indicates that these compounds have very different effects on chromatin, while the positive connectivity in P100 among all three compounds suggests that there are common signaling pathways that are activated regardless of whether sirtuins are activated or inhibited. Looking at GCP profiles, it was evident that resveratrol induced opposite effects from the other two compounds. Taken together, P100 data recognized that there was a common thread among the compounds while GCP data provided the finer details about their opposing mechanisms.

This first large-scale study of systematic perturbations with these proteomic readouts has clearly demonstrated the ability of the assays to capture cellular responses to a diverse set of therapeutic and investigational drugs. The proteomic assays perform on-par or better than other omics readouts where similar studies have been performed in terms of replicate reproducibility and MoA class detection. Subtle distinctions in MoA can be revealed through the sensitive nature of the assays to dynamic biochemical processes that may be independent of transcriptional programs. The high initial quality of the data in the resource suggests that proteomic assays are powerful tools to aid in drug characterization and are competitive with complementary technologies.

### Global analysis of connectivity profiles suggests that diverse signaling states converge to a restricted set of chromatin states

Our resource contains over 580,000 potential pairwise connections among 540 distinct drug-cell combinations. Connectivity analysis provides an objective measure of associations among perturbations in cells. We can define a *connectivity profile* for each perturbation: simply a vector of connectivity scores to all other compounds (including itself). Because they are quantitative, connectivity profiles can be analyzed in the same manner as raw assay profiles with techniques like clustering and principal component analysis. However, they have the advantage of reducing assay-specific considerations and provide a framework for comparing and integrating data across multiple assay types (see below).

We first sought to understand the structure of the connectivities as a whole. As a first step, we projected all within-cell connectivity profiles (allowing drug-drug connections only within a single cell type) using t-stochastic neighbor embedding (t-SNE; Figure 3A, Supplemental Data) (Maaten and Hinton, 2008). We noticed that both P100 and GCP connectivity profiles organized into spatial clusters based on cell type. This was not true when P100 or GCP *raw* profiles were projected in the same manner. There was far less structure in the projections, and what structure could be found correlated more with drug mechanism (Figure S3). This spatial clustering of connectivity by cell type supports the notion that individual cell types are “wired” differently. Drugs that connect in one cell type may not connect in another, and likewise for entire groups of drugs.

**Figure 3.**
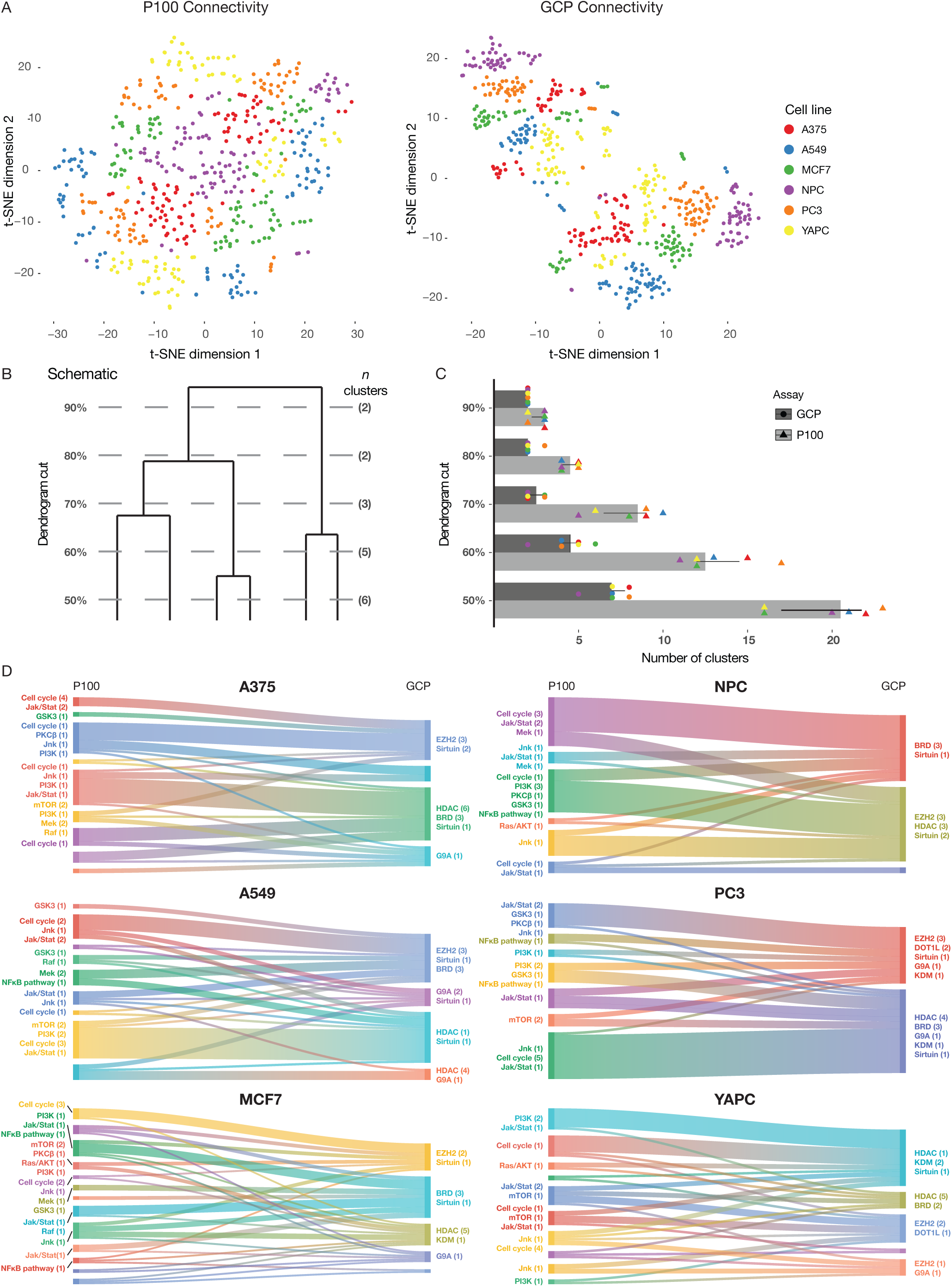
Global connectivity profile analysis. (A) t-SNE projection of P100 and GCP connectivity profiles for connections within each cell type (distance metric = Pearson correlation, perplexity = 60, learning rate = 10). (B) Schematic representation of cutting a dendrogram at fixed percentage of its height and counting resulting clusters, for illustration only. (C) Number of connectivity clusters formed as a result of cutting dendrograms as depicted in (B). Individual data points (six per assay) are overlaid on the boxplots and jittered on the y-axis for clarity. Error bars represent the 25th and 75th percentile limits. (D) Connectivity flows from P100 connectivity clusters to GCP connectivity clusters. For each cell line, only compounds reproducible in both assays are included. Each cluster is annotated by the major pertinent mechanistic classes for each assay, with the number of drugs in each class shown in parentheses. Colors are arbitrary. Because the analysis was restricted to the reproducible compounds in each cell type and single member clusters were eliminated, the number of clusters at the 60% cut for P100 and GCP may differ slightly from panel (C). See also Figure S3.

We further noticed that the projection of GCP connectivity seemed more highly structured than that of P100 (Figure 3A, compare right vs. left), with fewer and tighter spatial clusters, each encompassing more data points. This difference in the number of connectivity profile clusters led us to wonder whether there were more phosphosignaling states than chromatin states that could be adopted by individual cell types in general. In order to address this question, we hierarchically clustered drugs (1 - Pearson correlation) by their connectivity profiles in each cell type and cut the resulting dendrograms at fixed percentages of their height (depicted schematically in Figure 3B, results Figure 3C). This analysis demonstrated that the number of clusters grows more quickly for P100 connectivity profiles than GCP connectivity profiles as the cut percentage decreases. This alternative means of analysis provides further support that there are more phosphosignaling states available to cells than chromatin states.

To examine the cell type-specific properties of connectivity profile clusters, we mapped drugs from phosphosignaling connectivity clusters to chromatin connectivity clusters (analysis was limited to “reproducible” compounds in both assays in each cell type as identified in Figure 2, and dendrograms were cut at 60% of their height; single member clusters were eliminated). We counted the number of times a drug from a phosphosignaling cluster was present in a chromatin cluster, and generated connectivity “flow” diagrams that represent this mapping (Figure 3D). Phosphosignaling clusters were annotated by the major signaling-active drug mechanisms in the data set, and chromatin clusters by the major epigenetically-active drug mechanisms analogously. These flow diagrams showed diverse phosphosignaling states channeling into a smaller number of chromatin states in a cell type-specific manner. MCF7 and YAPC cell lines appeared to have the most complicated flow structure, while PC3 and NPC cells were the most simplistic. Frequently, signaling clusters containing cell cycle inhibitors mapped to chromatin clusters containing EZH2 inhibitors. Interestingly, NPC cells did not develop distinct chromatin connectivity clusters for EZH2 and HDAC inhibitors, although these classes may separate at a deeper dendrogram cut. BRD inhibitor-containing clusters also demonstrated unusual behavior, sometimes associating with HDAC inhibitors, other times associating with EZH2 inhibitors, and occasionally forming their own clusters. When we attempted to create an aggregated connectivity flow diagram across all cell types using the same methods, a very high number of connectivity clusters were formed (data not shown). We suspect that this is due to the intrinsic differences in the range of phosphosignaling and epigenetic states available to each cell type. Indeed, the connectivity flows visualized in Figure 3D paint a complex regulatory picture where drug perturbations have cell-type specific effects that are potentiated by available responses. The one unifying theme among all connectivity flows was that a variety of different signaling states can be induced in each cell type, and these coalesce to a relatively smaller number of chromatin states.

### Comparison of proteomic and transcriptomic connectivities demonstrates the strengths of assays measuring different biological dimensions

We envision that one major use of this resource will be to add value to, rather than replace, larger scale profiling efforts using other readouts. To illustrate how transcriptional and proteomic profiling can be utilized in tandem, we produced data for our entire set of perturbagens using the L1000 transcriptional profiling assay (Subramanian et al., 2017). L1000 raw data were processed using the standard L1000 computational pipeline. Next, similarities and connectivities among perturbations in L1000 were computed exactly the same way as for proteomic data using our universal connectivity framework (Figure 4A). In this way, we were able to directly compare connectivity results from proteomics (GCP and P100) to those from transcriptomics (L1000). Replicate reproducibility and connectivity within MoA classes in L1000 were similar to results observed for the proteomic assays (Figure S2).

**Figure 4.**
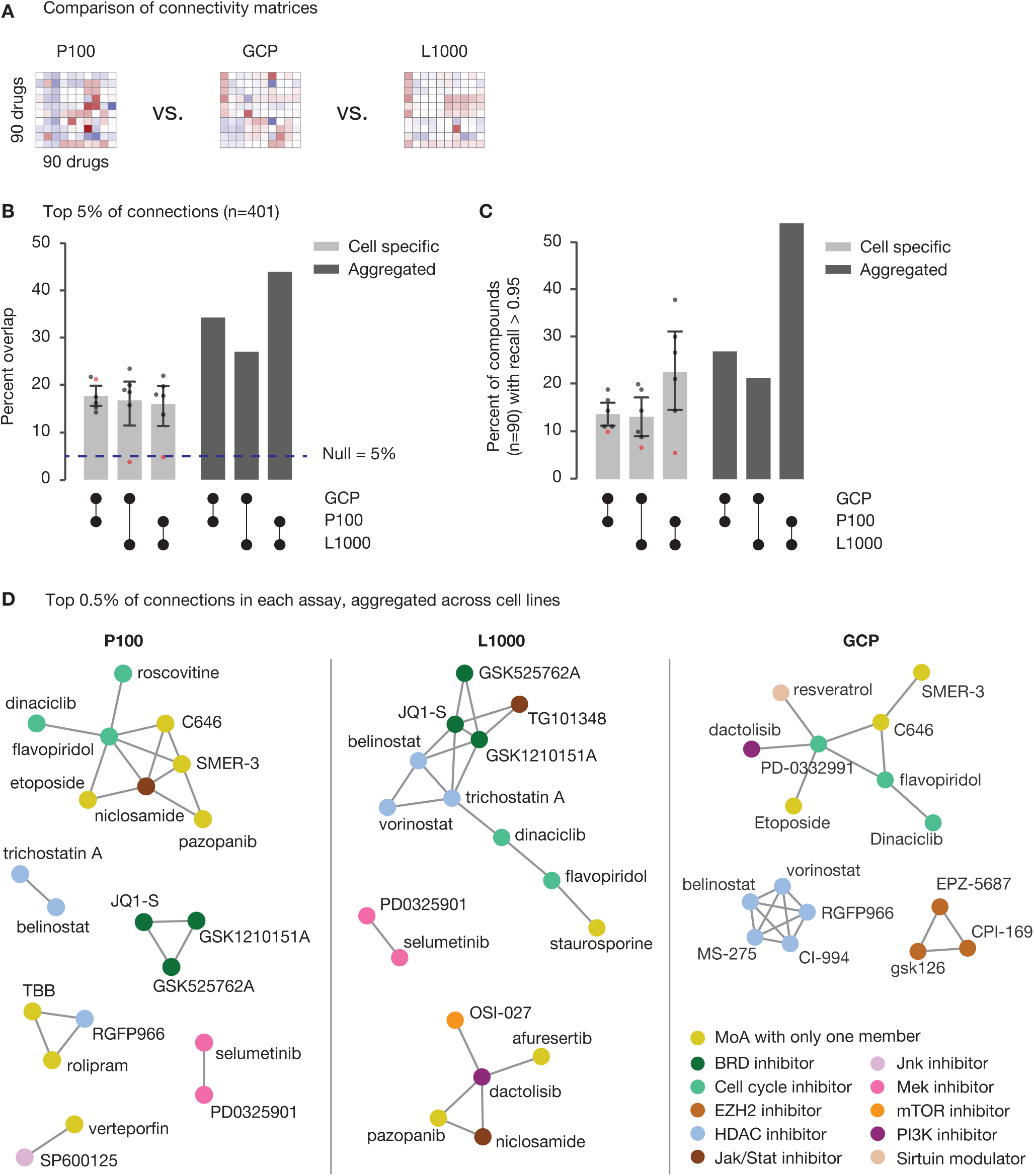
Comparison with transcriptomic data demonstrates assay-specific sensitivities. (A) Schematic of the comparison of connectivity matrices in three assays, including L1000 transcriptomic data. (B) Percent overlap of the top 5% of connections in the P100, GCP, and L1000 assays. The light gray bars shows percent overlap for cell-specific connectivities, and the dark gray bars show percent overlap for aggregated connectivities (median of six connectivity scores). The dashed line indicates the percent overlap expected by chance. (C) Recall of connectivity profiles across assays. The y-axis indicates the percent of compounds (n=90) that have recall greater than 0.95 for a pairwise comparison. Recall of 0.95 means that the connectivity profile for a particular compound in one assay had higher similarity to its corresponding connectivity profile in another assay than to 95% of other connectivity profiles (see Figure S4E for a schematic of this algorithm). Shading as in (B). (D) Network views of the top 0.5% of connections in each assay. All connectivity scores are positive. Compounds are represented by nodes, and MoA is encoded by the color of the node. See also Figure S4.

Our comparison of the three assays began with the following question: “How many of the strongest connections in one assay are also strong connections in another assay?” We defined a strong connection to be within the top 5% of connections for an individual cell line in each assay. We computed the percent overlap of the strongest connections (that is, the number of common connections divided by the number of considered connections) for each pair of assays either in each cell line individually or by aggregating all connectivities across cell lines using median (Figure 4B). We found that the top 5% of connections had an average of 16.8% overlap for cell-specific results and an average of 31.7% overlap for aggregated results. Therefore, aggregating connectivity scores nearly doubled percent overlap. This finding suggests that averaging over cell line-specific effects helps to improve agreement among the three methods of profiling. Both of these results are higher than the overlap expected by chance alone (5%; see Supplemental Note 3). Still, it is clear that different assays are most sensitive to different cellular responses, given that their strongest connections have little agreement, especially for cell-specific results. We repeated this analysis using 1% and 10% cutoffs with similar results (Figure S4A-B).

We also computed the percent overlap among all three assays concurrently. At the 5% cutoff, there was 6.0% overlap for cell-specific connectivities and 16.0% overlap for aggregated connectivities (Figure S4C). This percent overlap was lower than for pairwise comparisons (because it is harder for a strong connection to show up in all three assays compared to just two of them), but again, we observed that aggregation across cell lines improved overlap.

As an alternative method of comparing assays, we looked at whether individual drugs had similar patterns of connectivity across assays. In contrast to the previous analysis in which we considered the top connections in each cell line, this analysis treated drugs separately. For each drug, we compared its connectivity profile (see above) across assays (Figure 4C). We utilized an enrichment metric to compare connectivity profiles, but other similarity metrics, such as Spearman correlation, yielded similar results (Figure S4D). We found that an average of 14.8% of drugs had connectivity profiles that were highly similar across assays (recall > 0.95; see STAR Methods for a detailed description of the algorithm used). Briefly, recall of 0.95 indicates that a drug’s connectivity profile in one assay is more similar to its own connectivity profile in another assay than it is to 95% of the non-self connectivity profiles (Figure S4E). The percent of drugs with highly similar connectivity profiles more than doubled (33.7%) when connectivity scores were aggregated across cell lines. These quantifications of assay agreement were comparable to those reported by the overlap analysis (16.8% overlap without aggregation, 31.7% overlap with aggregation), indicating coherence between these two distinct analyses.

Both analyses highlighted NPCs as having especially low agreement across assays (red points in Figure 4B-C), yet replicate reproducibility was not appreciably lower in NPCs compared to the cancer cell models (Figure 2B). Our interpretation of these two findings taken together is that drugs treated in NPCs produced robust signatures, but many of these signatures were not distinct enough in the neural progenitor context to produce connectivity profiles that could be easily distinguished from the background. Figure 3D highlighted that NPCs have one of the least complex connectivity flows in the data set, and the restricted availability of signaling and chromatin states in NPCs is consistent with the integrative analysis here.

One novel observation revealed by the second analysis was that P100 and L1000 had considerably greater agreement with each other than with GCP. Considering aggregated connectivity scores, we observed that 53.3% of drugs had highly similar connectivity profiles (recall > 0.95) when comparing P100 to L1000, while that number dropped to 26.7% and 21.1% for comparisons to GCP. Considering that P100 measures the reduced phosphoproteome while L1000 measures the reduced transcriptome, it is not surprising that these two assays are more similar to each other than they are to GCP, which measures the more narrow readout of chromatin changes. The fact that this trend was not observed in the overlap analysis indicates that all three assays are capable of reporting on the strongest biological relationships, but there are individual compounds whose activity is not well captured through chromatin changes.

We sought to understand why there was little agreement among the strongest connections in each assay type. To investigate, we produced network views of the top connectivity scores for each assay aggregated (by median) across cell lines (Figure 4D), choosing only the top 0.5% of connections for visual clarity. These top 0.5% in each assay show examples of strong connectivity among compounds with mechanistic biases particularly likely to produce responses for the specific assay type. For instance, the GCP network shows five of the six HDAC inhibitors connecting strongly to each other, and all three EZH2 inhibitors connecting strongly to each other. We expected HDAC and EZH2 inhibitors to have strong signals in GCP because they are strong chromatin modifiers, and indeed we see connections between members of these classes in GCP, but not in the other 2 assays (at this stringent threshold).

A response to modulation of the cell cycle was captured by all three assays in slightly different ways. All three assays reported the connection of dinaciclib to flavopiridol, which are both cyclin-dependent kinase (CDK) inhibitors, but these compounds connected to a variety of other compounds in each assay. Our interpretation of these results is that different assays detect different components of cell cycle perturbation. It should also be noted that samples were collected at different time points post-treatment, possibly demonstrating a time-dependent evolution in response. This diversity of readouts could be beneficial for researchers interested in one particular type of cellular response; for example, a researcher seeking to discriminate subtle differences among a group of kinase inhibitors would likely be most interested in connectivity results reported by P100.

### Integration of proteomic and transcriptomic connectivities reveals cell line-selective vulnerabilities

In addition to serving as a complementary hypothesis generation tool, proteomic profiling will add value to transcriptomic profiling (and other profiling technologies) by providing reinforcing data. Especially for unexpected connections, we hypothesized that support from more than one assay would greatly increase a researcher’s confidence in the biological significance of a connection between two compounds. Therefore, we looked for examples of connections common to the three connectivity datasets.

To facilitate finding common connections, we computed the average connectivity score from the three assays; we refer to these data as AVG (Figure 5A). We visualized the strongest AVG connections aggregated across cell lines (again, by computing the median of six cell-specific connectivity scores) with a network view (Figure 5B). In contrast to the networks in Figure 4D, this network shows only connections with support from all three assays.

**Figure 5.**
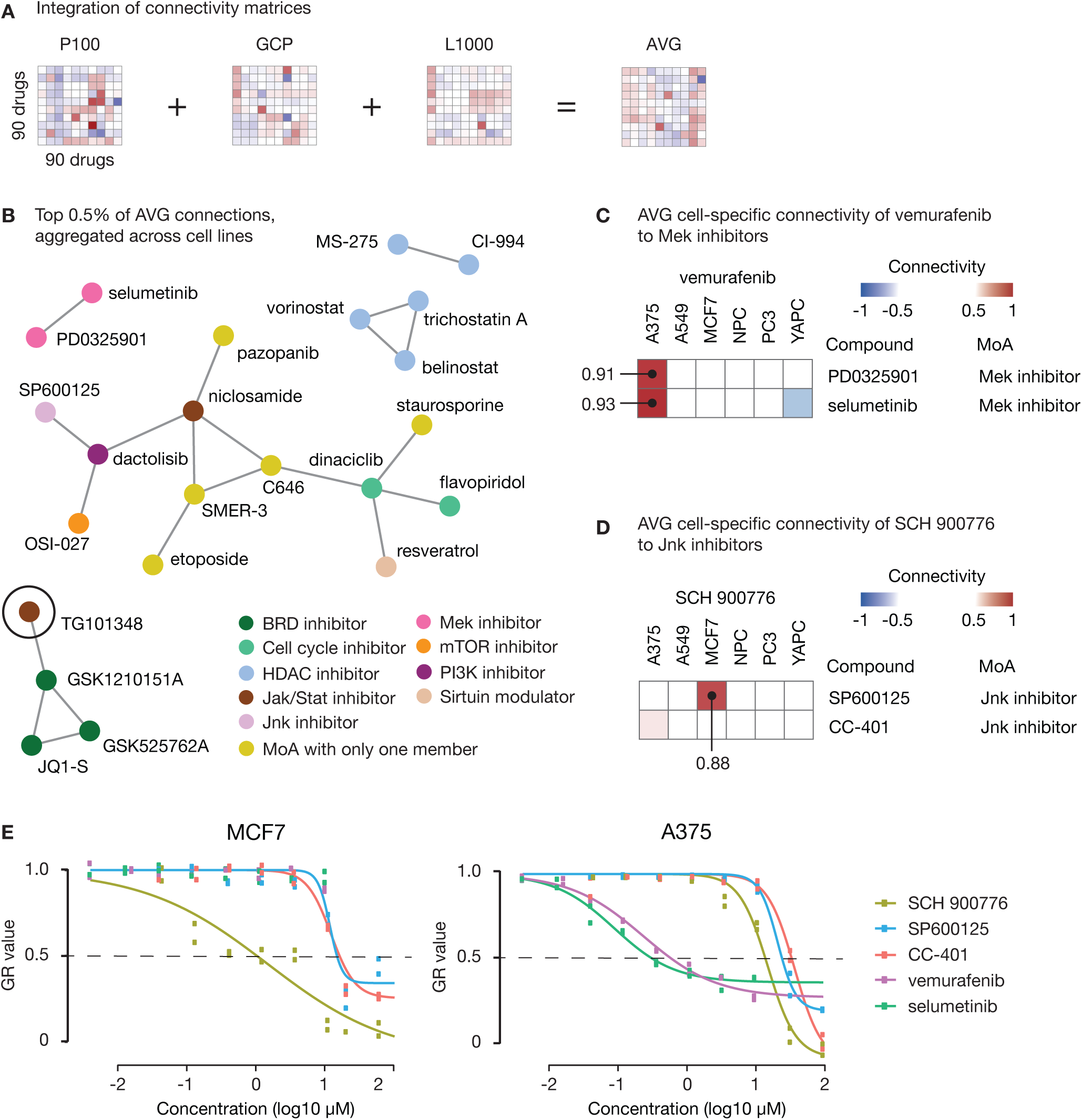
Multiassay data integration reveals cell-specific vulnerabilities. (A) Schematic of the integration of connectivity matrices in three assays to create AVG data. (B) Network view of the top 0.5% of connections for AVG data, which is an average of the connectivity scores in GCP, P100, and L1000. All connectivity scores are positive. Compounds are represented by nodes, and MoA is encoded by the color of the node. TG101348 (circled) has unexpected connectivity to the BRD inhibitors. (C) Heatmap view of the connectivity scores between vemurafenib and the two MEK inhibitors. The connectivity scores in A375 (0.91 and 0.93) are considerably higher than connectivity scores in any other cell line. (D) Heatmap view of the connectivity scores between SCH 900776 and the two JNK inhibitors. The connectivity score between SCH 900776 and SP600125 in MCF7 (0.88) is an unexpected cell-specific connection. (E) Results of a five-day follow-up viability experiment. The y-axis shows GR values in MCF7 (left) and A375 (right). GR values quantify drug cytotoxicity and are insensitive to different cell growth rates. The x-axis shows drug concentration on a log_10_ scale.

Apart from the large cluster containing two cell cycle inhibitors and a variety of other MoAs, all other clusters except one contained compounds with the same annotated MoA. The sole exception was the cluster containing TG101348 (circled) and the three BRD inhibitors. TG101348 is commonly annotated as a JAK2 inhibitor (Wernig et al., 2008), which makes its connection to BRD inhibitors perplexing. However, it was recently demonstrated through a BRD binding assay that TG101348 indeed has BRD inhibitor activity (Ciceri et al., 2014). This independent experiment strengthened our hypothesis that an unexpected connection seen by all three profiling assays would be likely to withstand experimental validation.

AVG data also reported compelling cell-specific connections. We immediately saw that vemurafenib (a BRAF inhibitor) was connected to the two MEK inhibitors only in A375 cells, an expected result. A375 cells are highly sensitive to both BRAF and MEK inhibition because they harbor the *BRAF*^*V600E*^ mutation that makes them dependent on RAF-MEK-ERK signaling (Wagle et al., 2011). Looking at cell-specific connections in AVG data, we confirmed that there was strong connectivity (0.91 and 0.93) between vemurafenib and the two MEK inhibitors in A375 cells and weak connectivity in all of the other cell lines (Figure 5C). The presence of these expected connections in all three assays indicates that inhibition of BRAF and inhibition of MEK appear similar to each other in all profiling modalities: chromatin, phosphoproteomic, and mRNA changes.

A cell-specific connection even more striking in its specificity than the previous example was that of SCH 900776 (SCH) to SP600125 (SP) in MCF7 (Figure 5D). SCH is a cell cycle inhibitor that targets CHEK1 (Bridges et al., 2016), while SP is a JNK inhibitor (discussed above). These compounds have different MoAs so it was unclear why they should connect, and it was especially unclear why they should connect in only one cell line. Noting the similarity of this connection to that of vemurafenib and MEK inhibitors in A375, which connect to each other in A375 because of differential cytotoxicity in A375 cells, we hypothesized that SCH connected to SP in MCF7 because these compounds were differentially cytotoxic in MCF7 cells.

We investigated our hypothesis with a follow-up viability experiment (Figure 5E). In order to account for variable cell growth rates, we quantified drug cytotoxicity using GR50 rather than IC50 (Hafner et al., 2016). The GR value represents cell count relative to DMSO control, adjusted for the cell line’s growth rate. We found that SCH was 13.8 times more cytotoxic in MCF7 (GR50: 1.03 □M) than in A375 (14.2 □M), and SP was 1.7 times more cytotoxic in MCF7 (GR50: 14.7 □M) than in A375 (GR50: 24.9 □M). For comparison, the GR50s of vemurafenib and selumetinib in A375 were 0.58 □M and 0.31 □M, which are comparable to the GR50 of SCH in MCF7 (1.03 □M). GR50s were not calculated for vemurafenib and selumetinib in MCF7 because 10 □M was insufficient to decrease the GR value below 0.5. These results confirmed our hypothesis that MCF7 was differentially sensitive to SCH and, to a lesser extent, SP. Furthermore, they suggest that these inhibitors might be candidate therapeutics for cancers similar to MCF7 in lineage (i.e. breast) or genetic features (e.g. estrogen receptor positive). Indeed, CHEK1 inhibition is a promising therapeutic direction in the treatment of breast and ovarian cancers (Bryant et al., 2014), and JNK signaling has an important, but poorly understood, role in breast cancer (Ashenden et al., 2017).

One modification to our original hypothesis is that CC-401, like SP, turned out to be more cytotoxic in MCF7 than in A375 cells. CC-401 was 2.1 times more cytotoxic in MCF7 (GR50: 16.1 □M) than in A375 (GR50: 34.4 □M). By our connectivity results alone, we might have predicted that only SP, and not CC-401, would be differentially sensitive in MCF7. One explanation for this discrepancy is that the dose of CC-401 (5 □M) was insufficient for perturbing cells in the manner that caused SP (25 □M) to connect to SCH. By including CC-401 in our follow-up experiment, we provided evidence that JNK inhibitors in general, rather than SP in particular, might be effective therapeutics for MCF7-like cancers. This anecdote highlights the caveat of any high-throughput hypothesis generation resource: while attempts are made to optimize as many experimental parameters as possible, follow-up experiments are indispensable for refining and validating therapeutic hypotheses.

### Proteomic connectivity links genetics to function and identifies potential therapeutic avenues

One powerful application of a library resource of systematic perturbation signatures is the ability to query existing, new, or even computationally derived phosphoproteomic or chromatin profiling data. To illustrate this application, we revisited data from a previously published chromatin profiling study that demonstrated how cancer-associated genetic alterations resulted in unique chromatin signatures (Jaffe et al., 2013). The Cancer Cell Line Encyclopedia is a collection of more than 900 diverse cancer lines representing different tissues and sites of origin and encompasses a wide variety of cancer driver mutations and downstream genetic dependencies (Barretina et al., 2012). In our previous work, we identified clusters of chromatin signatures related to gain-and loss-of-function of histone lysine methyltransferases. We set out to query these signatures against our library of systematic drug perturbations to correlate genetic features with drug activities.

To test this strategy, we first queried the chromatin profiles of cell lines belonging to a cluster identified as bearing EZH2 loss-of-function mutations in (Jaffe et al., 2013) (Figure 6A). EZH2 is a histone H3 lysine 27 (H3K27) methyltransferase capable of catalyzing the addition of methyl groups up to the fully saturated trimethyl state (Cao et al., 2002). Additionally, it is the core catalytic member of the Polycomb Repressive Complex 2 (PRC2), whose activity has been associated with repression of genes and marking of inactive chromatin (Kuzmichev et al., 2002). The signatures of the five cell lines in Figure 6A are characterized by the loss of H3K27me1, me2, and me3, with a concomitant increase in H3K27me0 and ac1. When these signatures are queried against our library of drug perturbations and ranked by their median connectivity scores, the top 18 positive connections are against EZH2 inhibitor compounds in the six distinct cell backgrounds present. This is, remarkably, the complete set of EZH2 inhibitor signatures present in the library.

**Figure 6.**
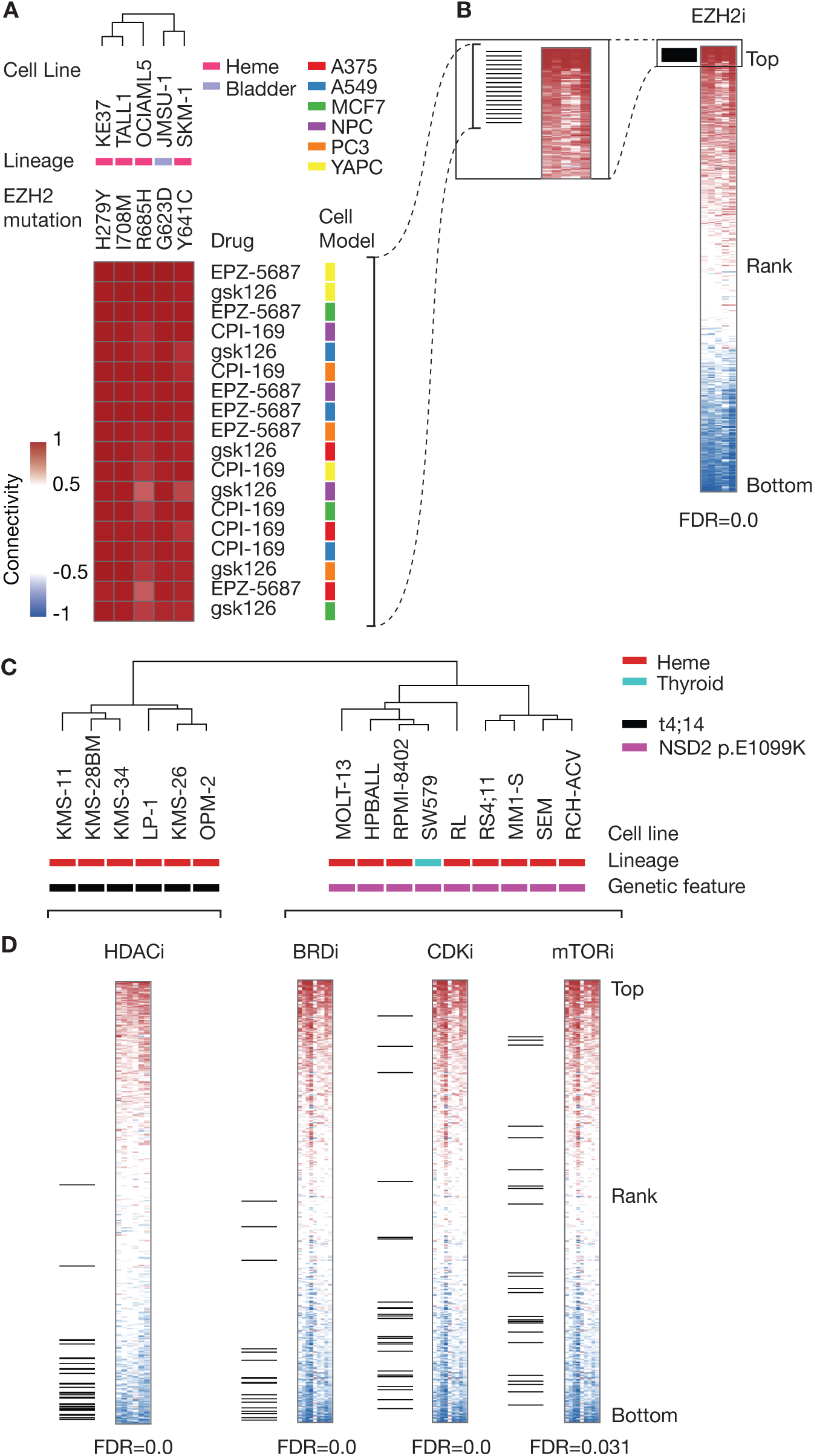
Connectivity query and perturbation set analysis of a diverse set of cancer lineages validates genetics and identifies potential therapeutic avenues. (A) Connectivity query of chromatin signatures from EZH2 loss-of-function cell lines from the CCLE. Results are sorted by the median connectivity to the perturbation across the five EZH2 loss-of-function cell lines. (B) Adaptation of the GSEA algorithm to test for enrichment of MoA classes in connectivity results. The top ranked set is EZH2 inhibitors with all hits to this set clustered at the top of the list sorted by average connectivity. (C) Stratification of two sets of NSD2 gain-of-function classes via hierarchical clustering of connectivity query results. (D) The most highly anti-connected perturbations to the *t4;14* and *NSD2*^*E1099K*^ gain-of-function classes of cell lines, when ranked by connectivity for each class. Enriched perturbation sets with FDR < 0.05 are shown for each class. The HDAC inhibitors are the most anti-connected perturbations to the *t4;14* subtype while BRD, CDK, and mTOR inhibitors are all anti-connected to the *NSD2*^*E1099K*^ subtype.

We adapted the technique of single sample gene set enrichment analysis (ssGSEA, or GSEA-preranked) to look for enrichment of MoAs in the query results (Barbie et al., 2009; Subramanian et al., 2005). Using our previously assigned MoA annotations, we tested for enrichment of each MoA in the rank-ordered connectivity query results. As a proof of concept, the most highly enriched MoA class for connections to EZH2 loss-of-function mutant lines was the EZH2 inhibitor class (Fig 6B, FDR=0.0).

Another key finding in (Jaffe et al., 2013) was the recognition that two classes of genetic events — *t4;14* translocation and *NSD2*^*E1099K*^ mutation — led to strikingly similar chromatin phenotypes based on the increase in H3K36 methylation levels (e.g. profiles from each class of genetic event did not resolve into distinct clusters but did segregate from other cell lines lacking these events). This observation led to the assignment of *NSD2*^*E1099K*^ as a gain-of-function mutation and a dependency in acute lymphoblastic leukemia (ALL) lines that harbored it. Building upon that previous work, we queried the chromatin signatures (obtained from a predecessor of the GCP assay) of *t4;14* and *NSD2*^*E1099K*^ cell lines from (Jaffe et al., 2013) against our resource of drug perturbation-induced chromatin signatures. Surprisingly, the connectivity profiles were able to perfectly segregate the *t4;14* and *NSD2*^*E1099K*^ classes of cell lines into different clusters (Figure 6C) despite the apparent similarity of the chromatin signatures that had been previously observed. This example illustrates that subtle differences in underlying signatures can drive vastly different connectivities and also demonstrates that using the entire profile for comparative analyses can provide more discriminatory power than simply analyzing differentially expressed analytes or typical clustering methods on raw profiles.

To further understand the basis for stratification, we again tested for enrichment of MoA classes in the connectivity query results. This time, we focused on the perturbations that were the most *negatively* connected to the signatures of the CCLE lines under the rationale that finding MoAs able to effectively reverse the chromatin signature induced by a genetic dependency might be a good means of selectively killing or sensitizing the cells (Figure 6D). Our enrichment analysis indicated that the signatures of the *t4;14* translocated lines were highly anti-connected to the HDAC inhibitor class of perturbations while the *NSD2*^*E1099K*^ lines were anti-connected to BRD inhibitors, CDK inhibitors, and mammalian target of rapamycin (mTOR) inhibitors in our data set (all at FDR < 0.05). These functional enrichments are highly consistent with investigational therapies for the specific cancer subtypes that typically harbor these mutation classes (see below).

The first class of genetic event, *t4;14* translocation, was composed entirely of multiple myeloma cell lines. The initial publication describing the CCLE (Barretina et al., 2012) included drug sensitivity for five of the six *t4;14* lines described here (KMS-28BM was not profiled) along with ~50% of all cell lines in the collection. One of the drugs tested in that study was panobinostat, a potent HDAC inhibitor. The panobinostat EC_50_ was in the 12th percentile or better for all of the *t4;14* multiple myeloma lines tested, and notably the EC_50_ of panobinostat against KMS26 was less than the first percentile at 3.7 nM. The insight provided by the negative connection of these lines to HDAC inhibitors is further supported by the fact that HDAC inhibitors have proven to be potent elements in the treatment of multiple myeloma, especially in combination with other agents (Laubach et al., 2017; Steiner and Manasanch, 2017).

The second class, *NSD2*^*E1099K*^, was composed of mostly acute lymphoblastic leukemia cell lines (a mixture of B-ALL and T-ALL, with the exceptions of SW579, a thyroid cancer line, and RL, a lymphoma line). Unfortunately, most of these lines were not profiled for drug sensitivity in the original CCLE study. However, several studies have recently emerged demonstrating the potential of mTOR inhibitors as therapeutics for ALL, especially in synergy with other agents (Iacovelli et al., 2015; Shi et al., 2016; Tasian et al., 2017; Witzig et al., 2015). One recent study details the synergy between an mTOR inhibitor and a CDK4/6 inhibitor in T-ALL, hitting two of the three classes of compounds that are anti-connected to the *NSD2*^*E1099K*^ set of lines. BRD inhibitors have also demonstrated therapeutic potential in B-ALL (Ott et al., 2012). The overlap among the mechanisms nominated by the proteomic connectivity analysis and these ongoing clinical studies is encouraging for the prospect of translational use of the proteomic resource.

Taken together, this evidence suggests that connectivity analysis might be helpful in suggesting potential therapeutic strategies based on anti-connections between genetic alterations and classes of drugs. It is not implausible that, if a disease induces a certain molecular state in a cell, then a compound that drives the cell towards an opposite state might help contribute to a resolution of the disease state. Our literature analysis suggests that this “driving” might be most helpful as a potentiator of other therapeutics or a means of overcoming resistance.

## Discussion

Increasingly, the improvement of profiling technologies coupled with high-throughput techniques has allowed for construction of systematic resources characterizing cellular responses to drug perturbation (Iorio et al., 2016; Rohban et al., 2017; Subramanian et al., 2017). Historically, proteomic techniques have been difficult to apply in the generation of such resources due to the heterogeneous sampling nature of mass spectrometry-based techniques as well as the cost and time required to generate the data. Yet proteins and dynamic changes thereto are important effectors in cells and are frequently the actual targets of drug therapies. Thus, it is valuable to create systematic resources with molecular readouts in the proteomic space to complement other techniques.

Here, we demonstrated the feasibility and value of creating a resource of drug perturbation proteomic signatures for signaling and epigenetics, two mechanistic classes being extensively explored for therapeutic development. Many landmark proteomic studies focus on the depth of coverage achieved at the expense of the number of samples profiled. Instead, we chose to focus on breadth rather than depth to generate the basis for an expandable resource. We created thousands of individual profiles of 90 different drugs in six different biological model systems. To overcome the time, costs, and stochasticity normally associated with deep discovery proteomics, we employed only compact, targeted assays that characterize high value analytes. We demonstrated that signatures from these assays are reproducible and that drugs in general are apt to produce strong signals in these molecular spaces (signaling and epigenetics) regardless of known or assumed mechanism of action. Because these assays have been reduced to practice, automated, and standardized, we can continue to build this resource over time and ensure that new data are easily comparable to those already collected.

There are other valuable precedents in proteomics of generating systematic resources. Li and colleagues have profiled more than 650 cell lines using reverse-phase protein arrays (Li et al., 2017), while Mertins and colleagues have performed deep proteomic characterization of over 100 breast cancer tumors (Mertins et al., 2016). Both of these efforts have focused on experiments of nature (genetic variation) rather than purposefully administered exogenous drug treatment. However, both studies demonstrated the value of a proteomic dimension in molecular profiling and concluded that it delivers complementary information to gene expression profiling. Here, we sought to extend the promise of the proteomic dimension in a direction that is suited to understanding drug mechanisms in cells, optimization of therapeutics under development, and nomination of therapeutic indications based on the induction of desirable proteomic signatures in cells (e.g. Figure 6).

One of our key motivations was to use assays that directly report on cellular biochemical processes likely to be directly affected by drug perturbation. To that end, we developed assays of phosphosignaling and chromatin biochemistry that directly observe the relevant molecules without need for antibodies. This allowed us to tailor the biochemical preparation and sample generation specifically for the desired analytes. One could fairly argue that space of the P100 phosphosignaling assay is relatively small compared to the number of known phosphosites in existence, and that not much is known about the biological function of specific analytes monitored in the assay. This makes it somewhat difficult to draw simple biological conclusions from the primary observations of changes in specific phosphopeptides (i.e. we don’t know much about the implications of the regulation of AHNAK phospho-S3426), as opposed to the GCP assay where there is general knowledge about repressive or activating tendencies of certain chromatin marks. However, the data-driven approach with which the analytes were selected has demonstrated strong reporting activity on a wide variety of drug perturbations. By intentionally neither selecting nor excluding sentinel markers from known pathways, we sought to avoid a bias towards reporting only on “known” activities. We also note that, while drugs and drug MoAs have strong replicate reproducibility in P100, changes in most individual sites themselves are relatively subtle, rarely exceeding two to three fold from median levels. Therefore, we believe that the relevant unit of comparison is the entire profile of a drug perturbation to that of *other* perturbations, rather than looking at the activity of specific markers within that profile.

Creating functional readouts in multiple biological activity spaces is a core goal of the LINCS program (http://lincsproject.org), under which this resource was developed. However, having distinct assays created a challenge of data integration. To address this challenge, we took a cue from earlier efforts (Lamb et al., 2006) and generalized the concept of connectivity — principled comparison of whole molecular profiles in large data sets with the goal of recognizing different biological conditions that induce related biochemical states in cells — to our proteomic data. Connectivity allows us to have a uniform way of asking the questions, “given molecular profile data induced by a drug treatment, what else does this look like? Have we seen something like this before?”

Importantly, our adaptation of connectivity is independent of assay type, which allows for facile integration of data from our discrete assays by asking whether the same connections are found in each (e.g. does the query of flavopiridol return the same connections in both P100 and GCP?). In this study, we used the exact same framework to further integrate transcriptional profiling data obtained for the same experimental conditions. We found it somewhat surprising that only a relatively small number of connections consistently rise to the top in all three activities assayed here (Figure 4). This finding supports the notion that there is no “one size fits all” approach to characterizing cellular responses (i.e. by only measuring transcriptional profiles), and that different assays provide unique windows into the responses to different classes of perturbagens. For example, our chromatin assay seems particularly well-suited to characterizing responses to compounds with epigenetic targets, while the signaling assay may be blind to effects of these drugs in some cases. Yet we do not discount the complementary information offered by assays that can perhaps unexpectedly provide strong evidence of connectivity despite not directly assaying a drug’s annotated mechanism of action (e.g. BRD inhibitors have stronger intra-class connectivity in the signaling space than they do in the epigenetic space, Figure 2). These observations underscore the need for multiple readout activities to be considered for truly understanding biological systems and their responses to new milieux.

At the same time, the repeated observation of the same connection(s) among all three assay types discussed here seems to attach a special importance to the result. There is a strong tendency in biological research to attempt to generalize results over all of biology, yet at the same time to seek model-specific methods of manipulating systems (e.g. labeling a drug as a JAK2 inhibitor, yet searching for indications where a JAK2 inhibitor will selectively kill cancer cells driven by aberrant activity in JAK2 signaling). The framework of connectivity can help to distinguish the cases of true universality of response (i.e. Figure 5B, where connections are aggregated across assays and cell types) compared to cell type-specific responses (Figure 5C-D). The cell type-specific agreement of connections over three assays re-discovers the mechanism of one of the great recent advances in targeted cancer therapeutics: that a *BRAF*^*V600E*^ mutation (present in A375) confers sensitivity to the drug vemurafenib by acting in a manner similar to disrupting MEK activity. The connection of this selective BRAF inhibitor to the MEK inhibitors was only observed in A375 (Figure 5C) despite the universality of the MEK inhibition response itself (Figure 5B). We hope that many more such examples can be discovered as we continue to expand our data set and further analyze the one we have at present.

We emphasize that this resource is the foundation for establishing a systematic repository of proteomic signatures that will be useful for drug characterization and more. Through LINCS, we are programmatically and institutionally committed to its maintenance and expansion. The resource is expandable in any number of dimensions; for example, by generating profiles for more drugs, including genetic perturbations and other stimuli, and profiling more cell types or models of biology (including tissue samples or patient-derived iPS models). All of these activities are already underway in our LINCS Proteomic Characterization Center for Signaling and Epigenetics.

We can also imagine other creative ways to increase the impact of the resource. Future developments in the assay technologies might allow for greater sample multiplexing, thus raising the rate of resource expansion. More comprehensive mass spectrometry technologies (such as DIA/SWATH) might readily expand the number of analytes covered in targeted proteomics assays, helping to answer questions about biological pathway modulation directly from primary data. And we could also envision harvesting data from other public studies that are likely to contain quantitative information on analytes covered by our assays (for example, (Bhanu et al., 2016; Kulej et al., 2015; Luense et al., 2016; Sidoli et al., 2015)) for direct comparison to and integration with our data. All analytes in our assays were selected, in part, for their relatively universal observability, making data harvesting a realistic possibility. One could also envision expanding the resource in the direction of new fit-for-purpose assay panels with complementary readouts that might be of high value for understanding the protein dynamics of disease including ubiquitination, arginine methylation and non-histone acetylation or methylation.

The data from this study are available in a variety of forms (Table 1). For mass spectrometrists, raw traces are available at Panorama Web (https://panoramaweb.org/labkey/LINCS.url). For the computationally-savvy user, all proteomic signatures at multiple levels of processing are available for download and computational manipulation via GEO (accession GSE101406). For all users, we have developed web apps that enable quick access to connectivity data (https://clue.io/proteomics) and developed tutorials to illustrate common use cases for interacting with the resource.

Finally, we envision collaborative interactions with users of this community resource — especially those interested in the characterization and development of therapeutics — influencing its future directions. We hope to have demonstrated the value of adding systematic proteomic studies to drug characterization and development efforts, and that the use of connectivity is a powerful abstraction that enables integration across data types. Ultimately, the value of this resource can only be validated by the discoveries and insights that it enables in the future, but we are enthusiastic about the potential applications of this resource at present.

## Supplemental Notes

### Supplemental Note 1

The choice of 3 hours for the P100 assay and 24 hours for the GCP assay is justified by biological considerations. Phosphosignaling (“signaling”) cascades can occur extremely quickly (e.g., initial EGF-induced signaling peaks just minutes after stimulation (Olsen et al., 2006)), while modifications to millions of histone molecules required to produce a detectable change in bulk chromatin states may require longer time spans and genome replication. We acknowledge that there is no perfect time for collection of all samples, and more focused studies can be executed on alternative time scales. These times were chosen to achieve the broadest utility for our initial resource. Second, our algorithm for choosing treatment concentrations was to utilize public drug metabolism and pharmacokinetics (DMPK) and absorption, distribution, metabolism, and excretion (ADME) data to select the reported bioavailable concentrations of the drugs in serum. In the absence of such data, we consulted literature for EC_50_/IC_50_ values or effective concentrations used in cellular studies. In the absence of prior knowledge or given conflicting evidence, we generally chose 1 M as a default concentration (Table S1).

### Supplemental Note 2

We suspected that part of the reason that compounds were more reproducible in P100 was that P100 has more features (96 analytes) than GCP (59 analytes after processing), so it is better able to resolve subtle differences in cell state. To investigate how much the larger feature space of P100 contributed to compound reproducibility, we randomly downsampled the P100 feature space to match that of GCP (i.e. 59 analytes) and reevaluted replicate reproducibility (Figure S2). We found that the mean difference between P100 and GCP reproducibility decreased from 11.6 to 4.0 compounds. Therefore, we conclude that the increased feature space of P100 explains much of the difference in reproducibility, and that these extra features encode information with discriminatory power.

### Supplemental Note 3

By chance alone, we expect 5% overlap: a random 5% of connections from one assay compared to a random 5% of connections from another assay would be expected to have an overlap of 0.25% (of all connections), which, when normalized to the portion of connections considered, is 0.25% / 5% = 0.05 or 5%.

## Supplemental Figure Legends

**Figure S1.**
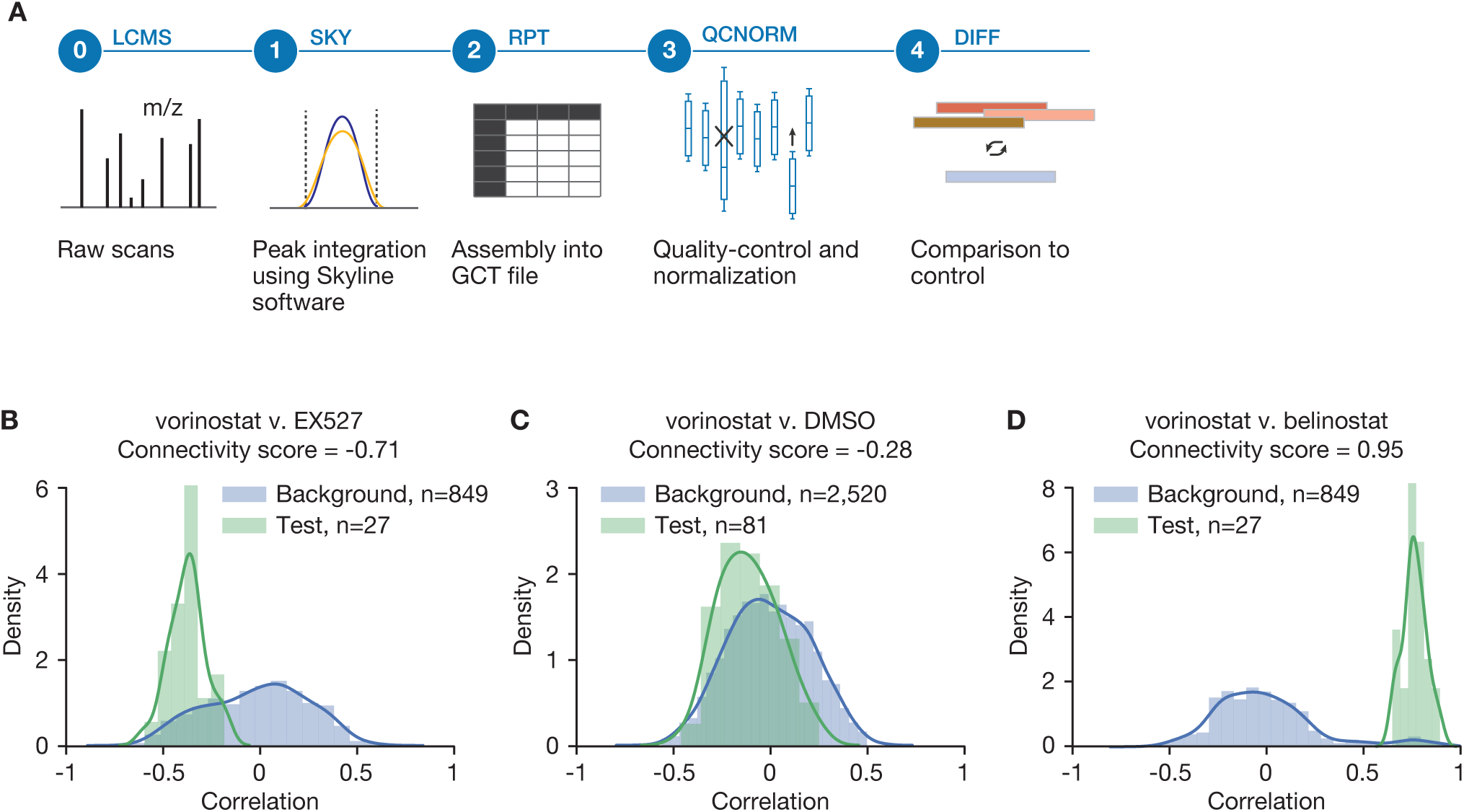
Data levels and examples of how connectivity is calculated. Related to Figure 1. (A) Schematic of the various data levels that are available. Data are processed in batches of 96-well plates. Level 0 is raw mass spectrometry (MS) scans. Level 1 is MS data that have been summarized by Skyline software to produce .sky files. Level 2 is log2 ratios of endogenous to internal standard peptides assembled into a GCT file. Level 3 has undergone quality-control and normalization. Level 4 is differential; that is, the signature for each sample is made relative to a control. The control can be either negative control wells, such as DMSO, on the same plate or all other samples on a plate. (B) Vorinostat v. EX527: example of a negative connectivity score close to −1. The background distribution (blue) consists of the correlations between the replicates of EX527 and all other samples. The test distribution (green) consists of the correlations between the replicates of EX527 and the replicates of vorinostat. (C) Vorinostat v. DMSO: example of a connectivity score close to 0. The background distribution (blue) consists of the correlations between the replicates of DMSO and all other samples. The test distribution (green) consists of the correlations between the replicates of DMSO and the replicates of vorinostat. (D) Vorinostat v. belinostat: example of a positive connectivity score close to 1. The background distribution (blue) consists of the correlations between the replicates of belinostat and all other samples. The test distribution (green) consists of the correlations between the replicates of belinostat and the replicates of vorinostat. Panels B-D show data for the GCP assay in A375 cells.

**Figure S2.**
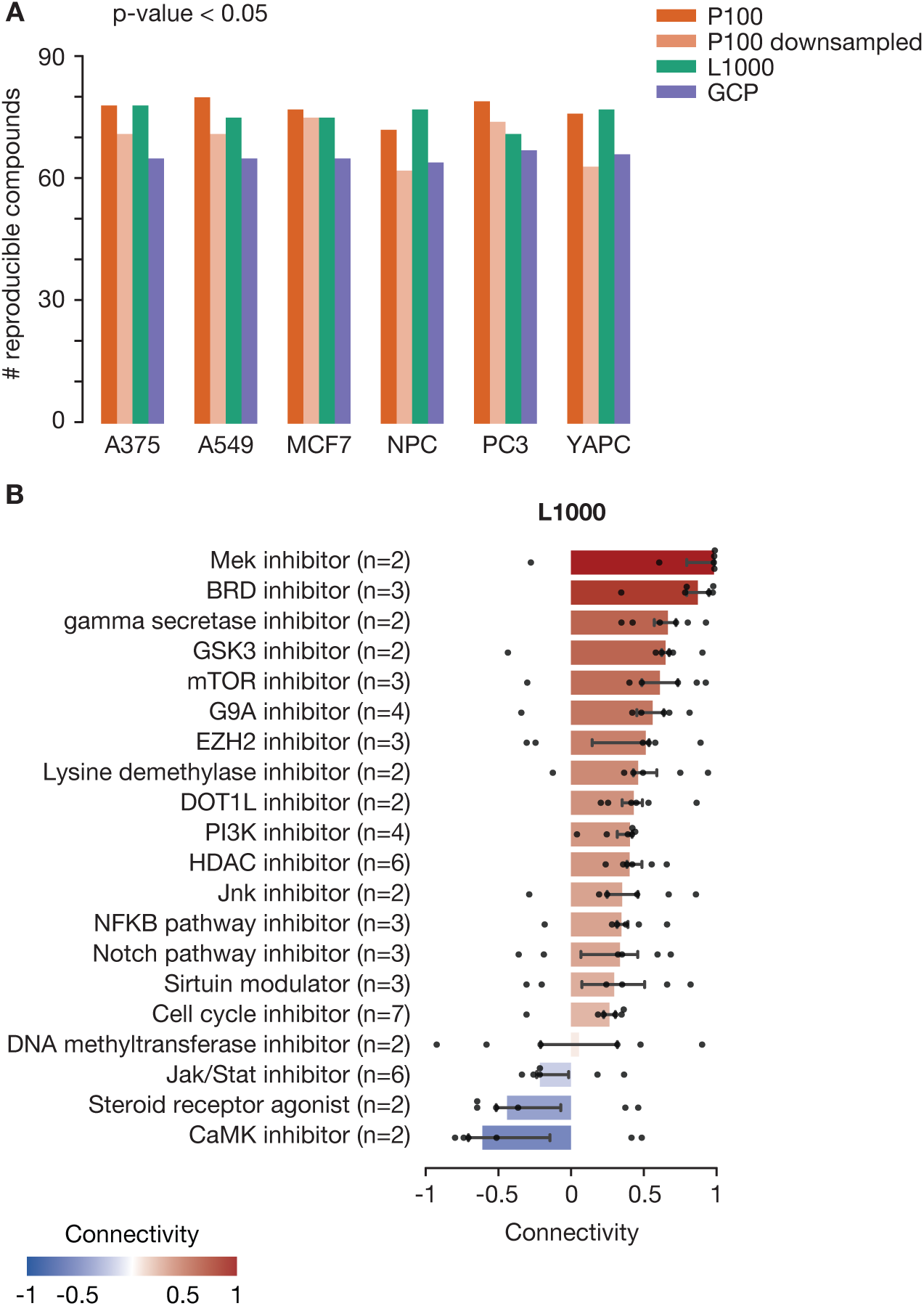
Replicate reproducibility for P100 downsampled and L1000, and MOA analysis for L1000. Related to Figure 2. (A) Bar chart showing the number of compounds considered reproducible in P100, P100 downsampled to have the same number of analytes as GCP (n=59), L1000, and GCP for each cell line. A compound was considered reproducible if the correlations among its replicates were significantly higher (p-value < 0.05) than the correlations among randomly chosen samples. Results for GCP and P100 have already been presented in Figure 2B and are shown here only for comparison. (B) Median connectivity of MoA classes in L1000. The extent of each bar is the median of six cell-specific median connectivities; error bars represent the 25th and 75th percentiles.

**Figure S3.**
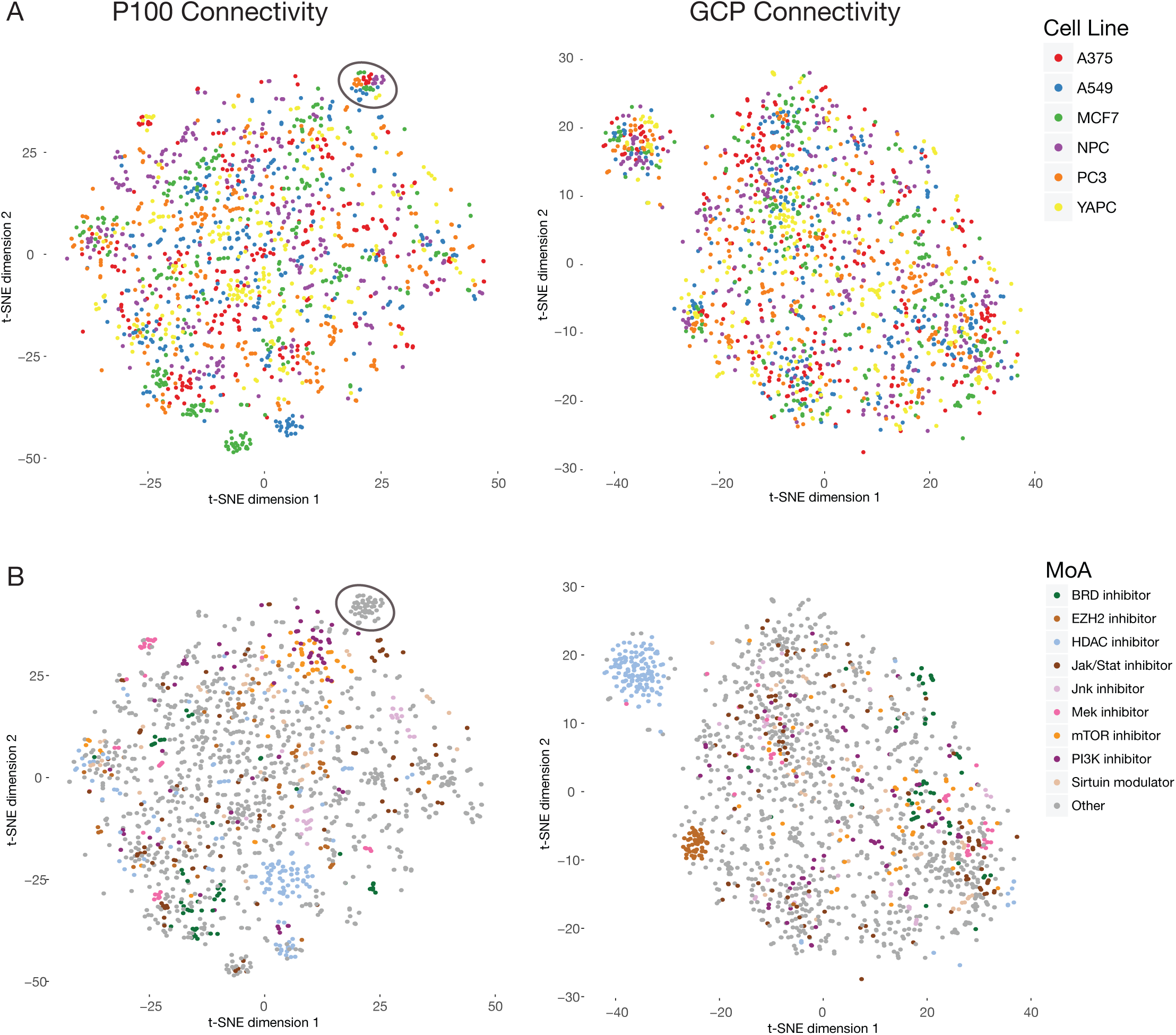
t-SNE of P100 and GCP signatures. Related to Figure 3. (A) t-SNE projection of P100 and GCP raw signatures (perplexity = 60, learning rate = 10), colored by cell type. The circled region in the upper right of the P100 plot corresponds to a cluster of staurosporine signatures. (B) Same t-SNE projection as in (A), except now colored by mechanism of action.

**Figure S4.**
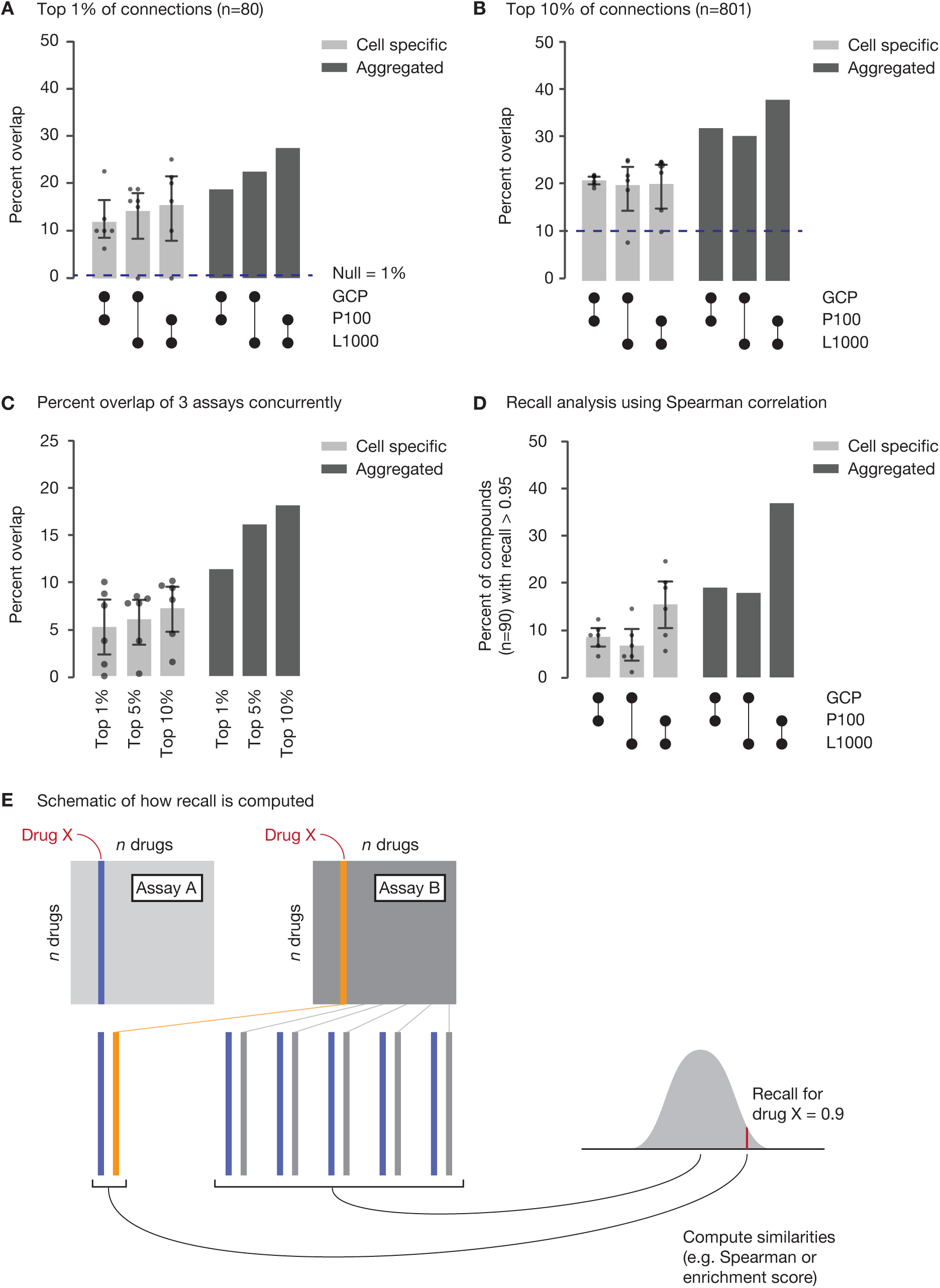
Alternative analyses comparing assays. Related to Figure 4. (A) Percent overlap of the top 1% of strongest connections (n=80) in the GCP, P100, and L1000 assays. The light gray bars shows percent overlap for cell-specific connectivities, and the dark gray bars show percent overlap for aggregated connectivities (median of six connectivity scores). The dashed line indicates the percent overlap expected by chance. (B) Percent overlap of the top 10% of strongest connections (n=801). (C) Percent overlap of three assays concurrently at 1%, 5%, and 10% thresholds. The percent overlap expected by chance depends on the percent of top connections considered; for the top 1%, 5%, and 10% of connections, the percent overlap expected by chance alone is, respectively, 0.01%, 0.25%, and 1%. (D) Recall analysis using Spearman correlation, rather than an enrichment metric. (E) Schematic illustrating how recall is computed. See STAR methods for a detailed description of this algorithm. Briefly; for a pairwise comparison of assays, recall for drug X measures how well that drug’s connectivity profile in one assay finds its matching connectivity profile in the second assay.

## Acknowledgements

The authors would like to gratefully acknowledge Dr. Anne Carpenter, Dr. Max Macaluso, Dr. Rajiv Narayan, Dr. Jodi Hirschman, and Bang Wong for extremely helpful discussion on the manuscript. We would also like to thank Dr. Maria Kost-Alimova for assistance with image-based cell viability assays. This work was funded in part by the NIH Common Fund’s Library of Integrated Network-based Cellular Signatures (LINCS) program by U54HG008097 (J.D.J.) and U54HG008699 (T.R.G. and A.S.)

## Author Contributions

Conceptualization, A.S., L.H.T., M.J.M, and J.D.J.; Methodology, L.L., R.P., J.G.A., A.L.C., J.D.E., C.M.D., T.K., T.E.N., J.P., and J.D.J.; Software, L.L. B.M., V.S., J.K.A., J.G., C.T., Z.L., and J.D.J.; Formal Analysis, L.L., R.P., D.L.L., and J.D.J.; Investigation, L.L., A.L.C., J.F.D., D.D., S.E., C.M.D., T.K., S.A.J., D.L., N.J.L., X.L., A.M., A.O., M.P., J.P.; Resources, L.H.T., M.J.M.; Data Curation, L.L., D.L.L., J.D.J.; Writing - Original Draft, L.L. and J.D.J.; Writing - Review & Editing, T.E.N., S.A.C., A.S.; Visualization, L.L., A.T., J.D.J.; Supervision, A.M., M.P., J.Z.Y., M.J.M., L.H.T., and J.D.J.; Project Administration, J.P., M.P. and J.D.J.; Funding Acquisition, T.R.G., A.S., and J.D.J.

## STAR METHODS

### CONTACT FOR REAGENT AND RESOURCE SHARING

Further information and requests for resources and reagents should be directed to and will be fulfilled by the Lead Contact, Jacob D. Jaffe.

### EXPERIMENTAL MODEL AND SUBJECT DETAILS

#### Cancer cell lines

After thawing from −80°C storage, cancer cell lines were recovered in standard tissue culture-treated dishes. Cells were allowed to propagate at 37°C and 5% CO_2_ for at least three doublings or until we had approximately 0.5 million cells per well of a six-well dish. A375 (female), A549 (male), and YAPC (male) cells were cultured in RPMI 1640 medium (Thermo Fisher). MCF7 cells (female) were cultured in DMEM (Thermo Fisher). PC3 cells (male) were cultured in RPMI 1640 medium containing 1 mM sodium pyruvate and 10 mM HEPES (Thermo Fisher).

DNA fingerprinting was used to authenticate the identity of cancer cell lines. Fingerprinting was performed at the Genomics Platform of the Broad Institute of MIT and Harvard (Cambridge, MA) using Fluidigm technology. The Fluidigm fingerprint panel includes a total of 96 SNPs, including 9 SNPs that overlap with the Affy 6.0 array and have multiple proxy SNPs each, 66 SNPs that overlap with Illumina’s 1m and 2.5m arrays and have multiple proxy SNPs each, 32 SNPs in transcribed regions of housekeeping genes that are expressed in most cell types, and 1 gender determining SNP.

#### Neural progenitor cells (NPCs)

Individual colonies of H9 human embryonic stem cells (ESCs) were cultured with mTeSR1 media (StemCell Technology) in matrigel (BD Biosciences)-coated plates. For NPC induction, ESC cell colonies of 60-80% confluence were incubated in a 1:1 mixture of N-2 and B-27-containing media (see below) supplemented with 1 μm dorsomorphin (Tocris Bioscience) and 10 μm SB 431542 (Tocris Bioscience). ESCs differentiated to NPCs in a single passage, and NPCs were cultured for nine passages.

N-2 medium consisted of DMEM/F-12 GlutaMAX (Thermo Fisher), 1x N-2 supplement (Thermo Fisher), 5 μg/ml insulin (Sigma), 1 mM L-Glutamine (Thermo Fisher), 100 μm MEM nonessential amino acids solution (Thermo Fisher), 100 μM 2-Mercaptoethanol (Sigma), 50 U/ml penicillin and 50 mg/ml streptomycin (Thermo Fisher).

B-27 medium consisted of neurobasal medium (Thermo Fisher), 1x B-27 supplement (Thermo Fisher), 200 mM L-Glutamine (Thermo Fisher), 50 U/ml penicillin and 50 mg/ml streptomycin (Thermo Fisher).

### METHOD DETAILS

#### Drug treatment

Cells were plated into six-well dishes 24 hours before treatment with one of 90 drugs from Table S1. Media was changed the morning of treatment. Cells were treated for either 3 (P100), 6 (L1000), or 24 (GCP) hours at 37°C before undergoing one of the following assay protocols. For each drug, at least three biological replicates were performed.

#### P100 assay

Please see Abelin *et al.* for a detailed description of the P100 assay (Abelin et al., 2016). Briefly; drug-treated cells were washed with cold PBS, lysed in-plate with urea buffer, and harvested by scraping. Cell lysates were transferred to a 96-well plate, flash-frozen using liquid nitrogen, and stored at −80°C until further processing. Upon thawing, samples (500 μg) were reduced, alkylated, and digested overnight with trypsin using the BRAVO Automated Liquid Handling Platform (Agilent). Peptides were desalted using a C18 Sep-Pak Cartridge (Waters) prior to immobilized metal affinity chromatography phosphopeptide enrichment. Phosphopeptides were enriched using commercially available Fe-NTA AssayMAP cartridges (Agilent). Salts were removed in a final desalting step using RPS cartridges (Agilent). A mix of isotopically labeled synthetic peptides was added to each sample prior to MS analysis. Peptides were separated on a C18 column (EASY-nLC 1000, Thermo Scientific) and subsequently analyzed by mass spectrometry (MS) as described in Abelin *et al*, or in DIA mode (Q Exactive^TM^-HF Orbitrap^TM^, Thermo Scientific). In DIA, full scans were acquired in the 300-1200 *m/z* range at 60,000 FWHM resolving power followed by DIA scans spanning *m/z* 400-1000 at 30,000 FWHM resolving power, using a 22 m/z isolation window and a NCE of 27. Alternating traversals of the DIA *m/z* range had their center isolation *m/z*s offset by 50%. Overlapping DIA scans were deconvolved with Skyline (MacLean et al., 2010) and analyzed exactly as in Abelin *et al*.

#### GCP assay

Please see Creech *et al*. for a detailed description of the GCP assay (Creech et al., 2015). Briefly; drug-treated cells were collected upon centrifugation. Upon lysis of the cells, histones were extracted using sulfuric acid and were precipitated using trichloroacetic acid. Samples (10 ug) were propionylated, desalted, and digested overnight with trypsin. A second round of propionylation was employed and samples were subsequently desalted using C18 Sep-Pak Cartridge (Waters). A mix of isotopically labeled synthetic peptides was added to each sample prior to MS analysis. Peptides were separated on a C18 column (EASY-nLC 1000, Thermo Scientific) and analyzed by MS in a PRM mode (Q Exactive^TM^-plus, Thermo Scientific) as described in Creech *et al*.

#### L1000 assay

Please see Subramanian *et al*. for a detailed description of the L1000 assay (Peck et al., 2006). Briefly; drug-treated cells were lysed using TCL Buffer (Qiagen), and mRNA transcripts were captured on oligo-dT-coated plates. Transcripts underwent ligation-mediated amplification (LMA); that is, mRNA was reverse transcribed to cDNA, gene and bead-specific probes were annealed to the cDNA, and probes were ligated and amplified via PCR using biotinylated universal primers. The PCR amplicon was then hybridized to beads with complementary oligonucleotide barcodes. After hybridization, the biotinylated amplicon was stained with streptavidin-phycoerythrin and detected using a Luminex FlexMap 3D system. The intensity of bead fluorescence corresponds to mRNA transcript abundance. Data were computationally processed using the standard Connectivity Map pipeline (Subramanian et al., 2017).

#### Data processing for GCP and P100

Mass spectrometry data files were imported into Skyline software (MacLean et al., 2010) in order to extract and integrate MS2 signals for endogenous (light) and synthetic internal standard (heavy) peptides. After manual quality assurance, Skyline files (.sky) were uploaded to a Panorama server (Sharma et al., 2014), where raw quantitative data were automatically assembled into Gene Cluster Text (GCT) files. The GCT format enables storing metadata and data in the same file. For each analyte (P100: *n* = 96, GCP: *n* = 61), the log_2_ ratio of the intensity of the endogenous peptide to the intensity of the internal standard peptide is reported. Samples are processed in batches of 96-well plates.

These data needed to be further normalized to enable comparison of samples within and across plates. The computational pipeline that performed these data processing steps is known as the Proteomics Signature Pipeline (PSP), and it is available online at https://github.com/cmap/psp. Importantly, the pipeline is automatically executed by the Panorama server on uploaded Skyline documents so all data processing operations occur independently of human intervention, yielding a reproducible research pipeline. The pipeline is self-documenting by appending a record of processing operations as metadata to each sample. First, any analytes or samples with an excess of missing data (thresholds are included in the provenance code for each plate and consistent within an assay) were filtered out. Next, a constant offset was applied to each sample in order to bring all samples to the same range. For GCP, we measure an analyte that is known to be invariant (histone H3, positions 41-49), so we subtracted this measurement from each sample. For P100, we do not have an invariant analyte, so we computed an analytical offset for each sample that minimized the distance between sample medians on a single plate. Finally, we made our signatures differential by subtracting from each analyte its median across a plate. Therefore, each final value is the log2 ratio of endogenous to internal standard peptide, relative to the median analyte measurement across all samples on a 96-well plate.

We make our data publicly available at multiple levels of processing (Supplementary Figure S1A). Skyline files (Level 1) and unprocessed GCTs (Level 2) for each 96-well plate are available on Panorama Web (https://panoramaweb.org/labkey/LINCS.url). Aggregated Level 2, Level 3 (quality-controlled and normalized) and Level 4 (differential) data are available via the Gene Expression Omnibus (accession GSE101406).

#### Viability follow-up experiment, related to Figure 5

A375 and MCF7 cells were thawed from −80°C storage, recovered in standard tissue culture-treated dishes, and allowed to propagated for at least three doublings at 37°C and 5% CO_2_. Cells were then plated into 384-well dishes and, twenty four hours later, treated with DMSO or one of the following drugs: vemurafenib, selumetinib, SCH 900776, SP600125, or CC-401. We performed an eight-point dose series in duplicate, where the maximum dose was 10 μm for vemurafenib and selumetinib and 100 μm for the other three drugs. Cells were treated for five days at 37°C.

We quantified cell viability by imaging for total cell counts. Images of cells were acquired using an Opera Phenix High-Content Screening Instrument (PerkinElmer) at 10x magnification in confocal mode, using brightfield and digital-phase contrast channels. Harmony software (PerkinElmer) was used to identify cells based on digital-phase contrast images and to count total cell numbers within four fields of view. Growth Rate (GR) values were computed, and sigmoid curves were fit using the online GR calculator (Hafner et al., 2016). The GR value represents cell count relative to DMSO control, adjusted for the cell line’s growth rate. Curves that did not reach a GR value of 0.5 were excluded.

### QUANTIFICATION AND STATISTICAL ANALYSIS

#### Computing similarities and connectivities

We utilize the framework of connectivity in order to compare perturbations across different profiling assays. We compute connectivity in two steps, and we do this separately for each cell line in each assay. First, we compute similarities among all profiles. We chose to use Spearman correlation, but other similarity metrics (e.g. Pearson correlation or Euclidean distance) could be substituted. Another similarity metric that we could have utilized is gene set enrichment analysis (GSEA), where one signature is compared to others by reducing it to up and down gene sets and asking for the enrichment of these gene sets in the other signatures. We chose to use correlation because we wanted to utilize *all* of our analytes. GSEA heavily weights the extreme tails of a signature, while Spearman correlation does not. Since correlation is sensitive to the length of vectors being correlated and our assays produce vectors (i.e. signatures) with different lengths, we cannot directly compare correlations across assays. For example, the correlations between L1000 signatures, whose lengths are 978, are drastically smaller than the correlations between GCP signatures, whose lengths are 61.

Therefore, we next convert correlations to *connectivity scores* by comparing observed correlations to a background distribution of correlations. The background distribution consists of all correlations between one of two drugs being compared and all other drugs profiled. Let us consider an example.

To compute the connectivity between two compounds A and B with three replicates each, we compare the correlations between the replicates of A and the replicates of B (test distribution; *n* = 9) to the correlations of the replicates of B with all other samples (background distribution; *n* ≈ 270). We use the two-sample Kolmogorov-Smirnov (KS) test to compare our two distributions. We opted for the non-parametric KS test because the assumptions of most parametric tests (e.g. normality of test distributions) are typically not satisfied by our distributions of correlation coefficients. The test statistic D of the KS test is what we call the connectivity score, except that we artificially add a sign to D in the following way: if the median of the test distribution is less than the median of the background distribution, the connectivity score becomes negative. Therefore, the range of connectivity scores is −1 (strong negative connection) to 1 (strong positive connection). When the test distribution consists of only one number (e.g. if we are comparing two drugs with only one replicate each), then the test statistic D simply becomes the fraction rank of a single correlation against the background distribution.

#### Global replicate v. non-replicate correlations, related to Figure 2A

We extracted all correlations among replicates, excluding the correlation between a sample and itself (because this value is always 1), and among non-replicates. We used kernel density estimation (KDE) to smooth these distributions.

#### Replicate reproducibility, related to Figure 2B

We sought to quantify whether the replicates of a drug were reproducible in order to determine whether that drug had a meaningful signal in a particular assay or cell line. For a given drug X with *k* replicates, we extracted the pairwise Spearman correlations 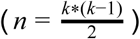 among its replicates and aggregated these values using median. Next, we created a randomly sampled null by picking *k* profiles at random from the ≈ 270 profiles in a given cell line-assay condition and computed the median of their Spearman correlations. We randomly sampled 1,000 times in order to create 1,000 medians in the null distribution. Each drug was assigned a p-value, which was defined as the fraction of values in the null distribution that were greater than the observed median correlation among replicates. For example, a p-value of 0.01 for a particular drug means that the replicates of that drug had a higher median correlation to each other than 99% of randomly selected, size-matched sets of samples from the same assay and cell line. In Figure 2B and Figure S2A, we considered drugs with p-values less than 0.05 to be reproducible.

#### MoA connectivity, related to Figure 2C

In each cell line, we computed the median of connectivities among compounds belonging to the same mechanism of action (MoA). Singleton MoAs were discarded. Each bar shows the six median connectivities (one for each cell line) for a particular MoA. The height of each bar is the median of these six values, and error bars represent the 25th and 75th percentiles of these six values.

#### Examples of MoA connectivity, related to Figure 2D

Each value in these matrices is a median of the six connectivity scores corresponding to the six cell lines.

#### t-SNE analysis, related to Figure 3A and Figure S3

Aggregated connectivity profiles (Supplemental Data, *n* = 540 per assay) were used to perform t-SNE analysis (Maaten and Hinton, 2008) in Morpheus (see below) using perplexity = 60 and learning rate = 10. Projections were rendered in R using ggplot2.

#### Clustering and dendrogram cutting, related to Figure 3C-D

A distance matrix was computed using the “correlation” metric of the Dist function in the R package “amap” for the connectivity profiles for each cell line (Supplemental Data, *n* = 90 per assay per cell type). This matrix was used to create dendrograms using the hclust function with “average” linkage. The function cutree was used to identify connectivity clusters at fixed percentages of the maximal heights of the resulting dendrogram. For Figure 3D, connectivity profiles were limited to the set of reproducible drugs (see “Replicate reproducibility” above) in each cell line, dendrograms were cut at 60% of their maximal height, and clusters with single members were eliminated. A network graph was constructed where nodes were considered individual connectivity clusters in P100 or GCP data, and edges were drawn between them with weight equal to the number of drugs shared between clusters. Network graphs were rendered using the “sankey” package of Google visualizations.

#### Percent overlap algorithm, related to Figure 4B

For each connectivity matrix, the top 5% of connections were extracted, except for self-connections. Self-connections were excluded since they are another way of quantifying replicate reproducibility, which was already addressed by previous analyses. Next, the percent overlap of these lists of connections were computed. For example, if the top 5% of connections in assay A, cell line C (*n* = 100) had 12 connections in common with the top 5% of connections in assay B, cell line C (*n* = 100), then the percent overlap was 12%. This computation was performed for each cell line separately as well as for a matrix in which cell lines were aggregated by computing the median of six cell-specific connectivity scores. For the cell-specific results, the height of the bar represents the mean, and the error bars show the 95% confidence intervals. This algorithm was repeated using 1% and 10% cutoffs for Figures S4A and S4B.

The null percent overlap corresponds to picking the top P% connections at random. For example, if *P* = 5, a random 5% of connections from one assay compared to a random 5% of connections from another assay would be expected to have an overlap of 0.25% (of all connections), which, when normalized to the portion of connections considered, is 0.25% / 5% = 0.05 or 5%. This estimate was computationally confirmed by repeating the random sampling many times in all cell lines.

#### Recall of connectivity profiles, related to Figure 4C

In order to assess whether a drug had the same pattern of connectivity in different assays, we compared *connectivity profiles* across assays. The connectivity profile for a given drug X is the vector of connectivity scores between drug X and all other drugs in a particular assay and cell line. We compared the connectivity profiles for the same drug in two different assays using Spearman correlation originally, but we found that considering only the tails of the connectivity profiles improved the similarity between connectivity profiles (Figure S4D). In order to look only at the tails of the connectivity profiles, we used a weighted enrichment score (Subramanian et al., 2005) instead of Spearman correlation. Finally, we asked whether the connectivity profile for drug X in assay A could recall its matching connectivity profile in assay B by computing a metric called *recall* (see Figure S4E). We defined the recall for a given drug X as the fraction rank of the similarity between matching connectivity profiles against similarities to all other connectivity profiles. For example, recall of 0.95 for drug X means that its connectivity profile in assay A, cell line C was more similar to its matching connectivity profile in assay B, cell line C than it was to 95% of all other connectivity profiles. A technical note is that the rank differs depending on which assay is considered first, so the recall for a particular drug is the mean of two ranks.

This computation was performed for each cell line separately as well as for a matrix in which cell lines were aggregated by computing the median of six cell-specific connectivity scores. For the cell-specific results, the height of the bar represents the mean, and the error bars show the 95% confidence intervals.

#### Querying external data against our resource, related to Figure 6

Using our framework of connectivity and the Proteomics Signature Pipeline (https://github.com/cmap/psp), we computed connectivity scores between all samples in our resource and the GCP profiles of 115 untreated cancer cell lines from the Cancer Cell Line Encyclopedia (CCLE) (Jaffe et al., 2013). This produced a matrix with 539 rows, representing 90 drugs in six cell lines (one drug had been excluded because of low technical quality), and 115 columns representing 115 cell lines from CCLE. For panel A, we extracted CCLE cell lines with *EZH2* mutations (*n* = 5) and sorted by the median connectivity score to these five cell lines. For panels B and D, we assessed enrichment of MoA classes in the 539 rows of drugs using GSEA (preranked by median connectivity across all sampled belonging to the genetic class: *n* = 5 for *EZH2*, *n* = 6 for *t4;14*, *n* = 9 for *NSD2*^*E1099K*^), but querying against compound sets rather than gene sets. Enrichment was assessed using the False Discovery Rate (FDR).

For panel C, we subsetted the connectivity matrix to cell lines with gain-of-function *NSD2* mutations (*n* = 15), either through *t4;14* translocation or the *NSD2*^*E1099K*^ point mutation. We clustered the columns of this subsetted matrix using Spearman correlation and the average linkage method. The two types of *NSD2* mutations segregated perfectly.

### DATA AND SOFTWARE AVAILABILITY1

#### Code for manipulating a GCT file

Software for manipulating a GCT file, which is the standard file format for the data discussed in this manuscript, is available in a variety of programming languages (i.e. Python, R, and Matlab). These repositories (cmapPy, cmapR, cmapM) can be found under the Connectivity Map team page on Github: https://github.com/cmap.

#### Proteomics Signature Pipeline

The software that processes proteomic data from Level 2 to Level 4, computes similarities and connectivities, and produces network visualizations of connectivity matrices is called the Proteomics Signature Pipeline (PSP). It is available at https://github.com/cmap/psp.

#### Proteomic Apps

In order to interact with the proteomic data discussed in this manuscript and additional data that will be released in the future, we have developed web applications for querying your own data against our resource and for exploring connections within our resource. The landing page for both of these apps is https://clue.io/proteomics.

#### Morpheus

Heatmaps were produced using a browser app called Morpheus (https://clue.io/morpheus).

#### Cytoscape

Networks were produced using Cytoscape software (http://cytoscape.org) (Shannon et al., 2003).

#### Data

All Level 2 through Level 4 data (for GCP, P100, and L1000) are available via the Gene Expression Omnibus (accession GSE101406). Level 1 data (Skyline files) and Level 2 data for individual 96-well plates are available on Panorama Web (https://panoramaweb.org/labkey/LINCS.url).

